# Single-cell transcriptomic landscapes reveal cell-type-specific regulatory mechanisms of nutrient accumulation and transport in wheat grain

**DOI:** 10.64898/2026.01.07.698077

**Authors:** Zhaoheng Zhang, Xiaohui Li, Xuelei Lin, Fuqiang Zhu, Qiangqiang Zhang, Jun Xiao, Yuan Chen

## Abstract

To elucidate the transcriptional regulatory mechanisms underlying wheat grain development, we constructed single-cell transcriptomic atlases of wheat grains at 4 and 8 days after pollination (DAP) using single-nucleus RNA sequencing. A total of 58,546 high-quality nuclei were clustered into 16 distinct cell populations representing all major seed tissues, including the embryo, aleurone layer, transfer cells, endosperm, inner pericarp, seed coat, and embryo surrounding region. Comparative analysis revealed a dynamic transcriptional reprogramming from growth-related pathways at DAP 4 to nutrient accumulation at DAP 8, which coincided with shifts in the expression of hormone signaling and organ development genes, as well as starch and storage protein genes. Pseudotime trajectory analysis identified two major differentiation branches, highlighting distinct roles in nutrient storage and support. Notably, the transcription factor *TaMADS58* was found to regulate pericarp cell number while simultaneously enhancing protein content and processing quality. Concurrently, *TaJEKLL*, specifically expressed in nucellar projection cells, was demonstrated to influence nucellar projection development and nutrient transport. Natural variation in *TaMADS58* and *TaJEKLL* revealed their potential to decouple yield from quality traits, offering promising applications for crop improvement. Collectively, this study provides a high-resolution single-cell resource for wheat seed biology and demonstrates that investigating gene expression with spatiotemporal precision at the single-cell level can facilitate the discovery of key genes for precisely regulating individual traits.

## Introduction

Wheat (*Triticum aestivum* L.) is one of the most widely cultivated crops worldwide and a major source of protein for the global population. Grain yield and quality are largely determined by endosperm development, which depends not only on starch and storage protein synthesis but also on the coordinated delivery of assimilates. During grain filling, wheat grain development and final yield depend not only on favorable environmental conditions but also on a precise and highly efficient internal nutrient transport system. The spike acts as the main source of assimilates, supplying sugars and amino acids directly to developing grains (Sanchez-Bragado et al., 2020). Within the seed, the nucellar projection, maternal pericarp, and transfer cells form a tightly coordinated conduit that channels nutrients into the endosperm (Wang and Fisher, 1994). This organized transport network ensures the timely delivery of assimilates to expanding storage tissues, supporting both endosperm growth and the accumulation of starch and storage proteins (Zhang et al., 2007).

Compared with maize and rice, wheat grains develop a distinctive ventral crease containing a specialized nucellar projection, which forms a narrow and structurally organized interface between maternal vascular tissues and the filial endosperm (Liu et al., 2022; Wang et al., 1994).Within this axis, transfer cells and adjacent endosperm regions are tightly juxtaposed, creating an efficient conduit for nutrient flow. *TaMADS29*, specifically expressed in the nucellar projection, contributes to maintaining the integrity of this tissue and supporting nutrient transport into the endosperm (Liu et al., 2023). During early grain development, the maternal pericarp undergoes dynamic remodeling, characterized by rapid expansion followed by progressive thinning and degradation, facilitating both the physical accommodation of the expanding endosperm and the directed transport of assimilates (Saada et al., 2021; Xiong et al., 2013). Starch granules in pericarp parenchyma cells undergo transformation and disintegration as the tissue degenerates, while endosperm transfer cells and starchy regions actively accumulate starch, indicating coordinated nutrient dynamics (Zheng et al., 2017; Zhuo et al., 2023). These coordinated anatomical and developmental processes establish a finely tuned nutrient transport network that integrates maternal and filial tissues, ensuring timely delivery of sugars, amino acids, and other assimilates to expanding storage tissues (Albacete et al., 2014; Patrick and Offler, 2001). Despite their importance for grain filling, how cells within the ventral crease and nucellar projection coordinate nutrient allocation remains unclear.

Building on this anatomical and functional framework, single-cell and single-nucleus RNA sequencing have emerged as paradigm-shifting approaches that enable the dissection of cellular heterogeneity, the reconstruction of developmental trajectories, and the inference of intercellular communication with unprecedented resolution (Wang et al., 2025; Yao et al., 2025). These technologies are particularly powerful for interrogating seed development, a process characterized by rapid cellular diversification and precise spatial coordination among maternal and filial tissues to support growth, nutrient allocation, storage and transport. Early work in Arabidopsis demonstrated that single-cell methods can capture endosperm transcriptional programs and elucidate the dynamics of genomic imprinting, underscoring their potential to reveal cell type–specific regulatory networks governing complex developmental processes (Picard et al., 2021). In major cereal crops, single-cell and spatial transcriptomic technologies have rapidly expanded our understanding of seed development. For example, in soybean, single-cell analyses of embryos and endosperm have delineated functional transitions among endosperm subtypes and identified sugar transporters linked to the transcription factor DOF11, offering a high-resolution perspective on yield- and quality-related regulatory networks (Zhang et al., 2025b). In maize, early endosperm differentiation establishes the major cell types that guide subsequent proliferation and nutrient accumulation. Single-cell RNA sequencing has revealed extensive endosperm heterogeneity, dynamic gene expression across cell types, and cell type–specific regulatory networks (Yuan et al., 2024),,while spatial transcriptomics has further resolved functional differences and mapped the spatial distribution of starch, protein, and oil (Fu et al., 2023). Integrated single-cell and spatial transcriptomic approaches therefore enable high-resolution mapping of cellular organization, developmental trajectories, and tissue-specific gene regulatory programs within complex plant organs. This integrative strategy has been well illustrated in rice, where combined analyses of germinating embryos resolved cell type – specific transcriptional dynamics underlying hormone signaling, starch metabolism, and coordinated inter-tissue nutrient transport (Yao et al., 2024). In wheat, spatial transcriptomic analyses spanning 4-12 days post-anthesis have provided the first comprehensive view of early grain development, revealing distinct spatial gene expression domains across the developing seed (Li et al., 2025).Building on this advance, we integrate single-nucleus RNA sequencing with spatial transcriptomics to generate a spatiotemporal, cell-resolved atlas of early wheat grain development across key stages following pollination. This delineates cellular heterogeneity, developmental dynamics, and tissue organization while preserving spatial context, enables the identification of cell type–specific regulatory programs, and clarifies how distinct tissues coordinate growth, nutrient allocation, and storage during early seed formation, providing a resource for dissecting complex developmental processes and informing targets for crop improvement.

## Results

### Single-cell transcriptomic atlas of early development stages in wheat grain

To elucidate the gene expression regulation at the cellular level during early wheat grain development, we generated single-nucleus RNA sequencing (snRNA-seq) libraries from wheat grains at 4 and 8 days after pollination (DAP) following manual removal of the pericarp, with two biological replicates for each stage (Fig. S1A). The average number of unique molecular identifiers (UMIs) per nucleus ranged from 2,000 to 10,000 (Fig. S1B), with 2,500-5,000 genes detected per cell (Fig. S1C). After stringent quality control, 58,546 high-quality nuclei were retained for further analysis (Fig. S1D). Unsupervised dimensionality reduction and clustering of single-nucleus transcriptomic data identified 16 distinct cell clusters (Fig. 1A). Each cluster ranged from 301 to 7,631 cells (Fig. S1E), reflecting the heterogeneity across seed tissues. Cluster 16 exhibited higher average UMI counts and expressed gene numbers than other clusters (Figs. S1F and S1G). Cluster-specific marker genes are shown in Figs. S1H and 1B, and their spatial distributions were annotated based on published wheat seed spatial transcriptomic data (Li *et al*., 2025) (Fig. 1C). We annotated all major cell types present in early wheat seed development, including the embryo, aleurone layer (AL), embryo-surrounding region (ESR), central cell of starch endosperm (CSE), transfer cells of starch endosperm (TCSE), Inner pericarp (IP), seed coat (SC), cavity fluid (CF), and Scutellum (Fig. 1B). For example, the *GDSL-type esterase/lipase* gene *TaLTL1-A1* is specifically expressed in the aleurone layer; *Alpha-amylase isozyme 3E* gene (*TaAMY3-A1*) shows specific expression in ESR cells; *Sucrose synthase 4* gene (*TaSUS4-D1*) is specifically expressed in the central endosperm; the *lipid transfer protein* gene *TaLTPL36-D1* is specifically expressed in TCSE cells; *Metallothionein gene* (*TaMT2b-D1*) exhibits specific expression in the IP; *ribulose-1,5-bisphosphate carboxylase small subunit* gene (*TaRBCS5-D1*) is specifically expressed in the SC; the *sugar transporter* gene *TaSWEET9-B1* is specifically expressed in the CF, *Nuclear transcription factor Y subunit B* gene (*TaNF-YB6-A1*) is specifically expressed in the Embryo, and the *Metallothionein* gene (*TaMT3a-D1*) high expression in Scutellum (Yao *et al*., 2024) (Fig. 1B, 1C, S1H, S1I, S1J and Supplemental Data Set S1). In addition, two clusters (0 and 16) highly enriched in S-phase and G2/M-phase markers were annotated as actively dividing cells (DC) (Fig. S1K and Supplemental Data Set S2). All clusters, except the embryo, exhibited higher inter-cluster correlations, highlighting the developmental distinctiveness of the embryo (Fig. S1L). We further performed pairwise comparisons between endosperm cell clusters identified in maize and wheat scRNA-seq/snRNA-seq datasets (Fig. 1D). The TCSE, AL, CSE, and ESR cells showed high marker gene similarity between maize and wheat (Supplemental Data Set S3) (Yuan *et al*., 2024). In addition, a large proportion of DC undergoing vigorous division corresponded to various maize endosperm cell types, suggesting their potential to differentiate into these tissues.

**Fig. 1.**
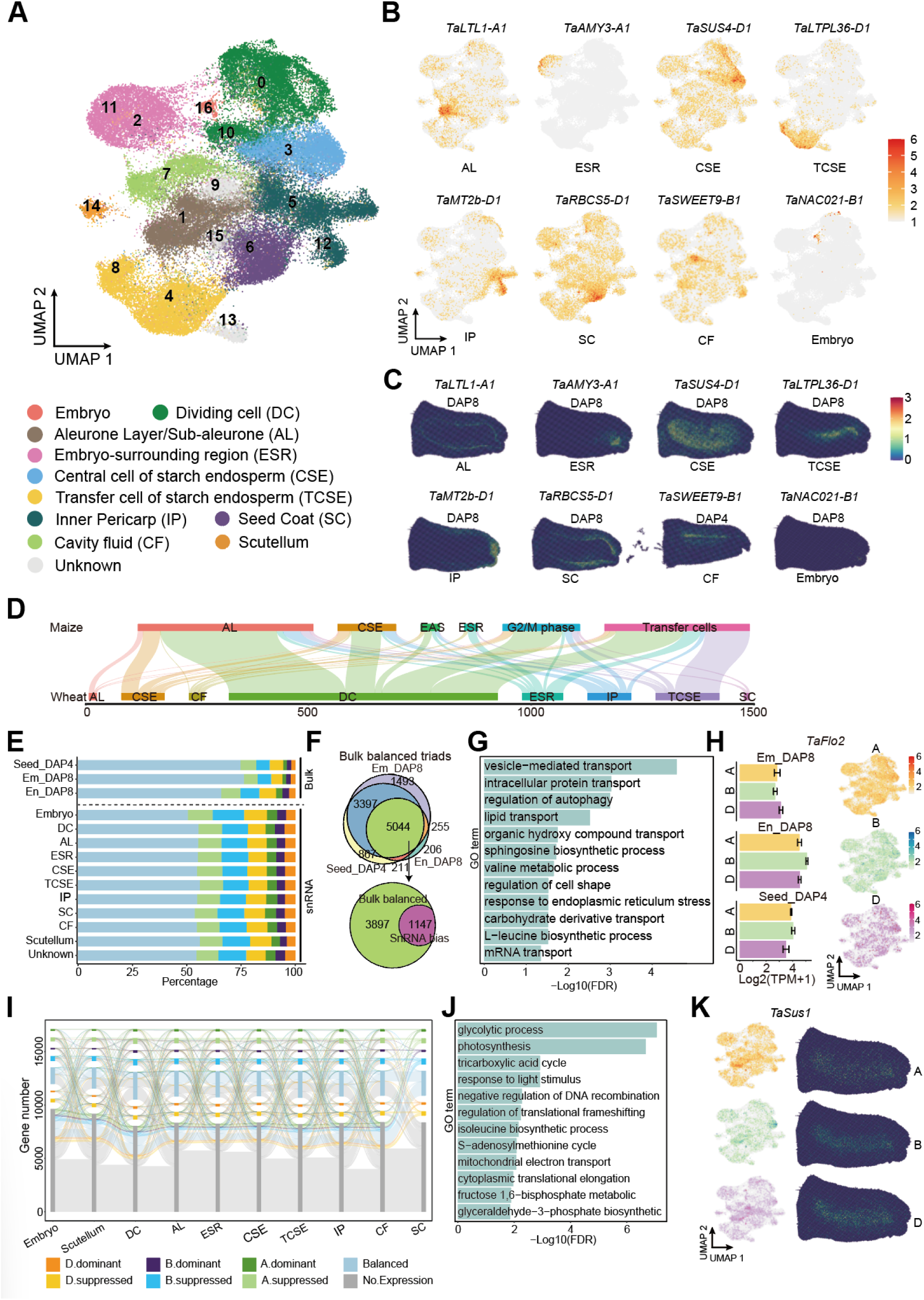
Construction of a single-cell transcriptomic atlas for developing wheat grain. A. UMAP visualization of wheat seed cells displaying 16 putative clusters and cell-type annotation. Each dot represents a single cell, color-coded by cell-types. B. UMAP plots of marker genes revealing the identities of cell-types. Each dot represents a single cell, color-coded by expression level. Color scale represents normalized expression levels. C. Spatial expression patterns of marker genes confirming cell-type identities. Color-coded by expression level. D. Sankey diagram showing that wheat shares degree of similarities in AL. CSE, CF, DC, ESR, IP, TCSE and SC cells with maize endosperm cells. The number of overlapped orthologous genes for each cell type is given on the bottom. E. Proportions of subgenome-biased genes detected in bulk RNA-seq compared with each cell-type in snRNA-seq. F. Venn plot shown the expression bias in snRNAseq of expression balanced triads from bulk-RNAseq G. GO terms enrichment analysis of 1147 genes from Fig.1F. *P*-values were obtained using hypergeometric test. The Benjamini-Hochberg (BH) were used to adjust the *P*-value. H. Expression levels of of the A-, B-, and D-subgenome homoeologs of *TaFlo2* in bulk-RNAseq and snRNAseq. Color scale represents normalized expression levels. I. Sankey diagram illustrating the transitions of subgenome-biased genes across cell-types. J. GO terms enrichment analysis of 488 triads that exhibited divergent expression bias in at least two tissues. *P*-values were obtained using hypergeometric test. The Benjamini-Hochberg (BH) were used to adjust the *P*-value. K. UMAP showing the cell cluster expression bias patterns of the A-, B-, and D-subgenome homoeologs of *TaSus1*. Color scale represents normalized expression levels.

Previous studies based on bulk RNA-seq datasets have reported that approximately 30% of wheat homoeologous gene triads exhibit asymmetric expression patterns (Zhao et al., 2023). To resolve homoeolog expression bias at higher resolution, we compared our snRNA-seq data with previously published bulk RNA-seq datasets derived from wheat embryos and endosperm (Zhao et al., 2024; Zhao *et al*., 2023). Notably, the snRNA-seq dataset identified substantially more genes with expression bias than bulk RNA-seq. Among 5,044 genes that displayed balanced expression across all three bulk RNA-seq datasets, 1,147 genes still exhibited subgenome-biased expression in distinct cell clusters in the snRNA-seq data, indicating that cellular heterogeneity in bulk samples masks cell-type-specific subgenome expression patterns (Figs. 1E, 1F and Supplemental Data Set S4). These 1,147 genes showing subgenome bias exclusively at the single-cell level were predominantly enriched in pathways related to nutrient transport and essential biosynthetic processes (Fig. 1G), suggesting their involvement in grain development and nutrient accumulation. For example, the grain-weight regulator *TaFlo2* exhibited pronounced repression of the D-subgenome homoeolog in snRNA-seq data, whereas no obvious subgenome bias was detected in bulk RNA-seq datasets (Sajjad et al., 2017) (Fig. 1H). In addition, Sankey diagram analysis revealed low similarity in homoeolog expression-bias patterns across different tissues, indicating that wheat homoeologous triads contribute to tissue-specific functional differentiation through distinct, tissue-dependent subgenome expression profiles (Fig. 1I). We further identified 488 triads that exhibited divergent expression bias in at least two tissues; these genes were mainly enriched in pathways associated with photosynthesis, glucose and derivative metabolism, and fundamental cellular processes (Fig. 1J and Supplemental Data Set S4). For instance, the A- and D-subgenome homoeologs of *sucrose synthase 1* gene (*TaSus1*) were highly expressed in AL cells, whereas the B-subgenome homoeolog was predominantly expressed in (CSE cells, indicating functional divergence of *TaSus1-B1* (Fig. 1K) (Wamalwa et al., 2020). Notably, such subgenome-specific expression patterns were not detected in spatial transcriptomic data. These results demonstrate that snRNA-seq captures cell type–resolved subgenome expression differences that are obscured in bulk and spatial transcriptomic datasets, providing a framework for interrogating subgenome functional divergence during wheat seed development.

### Transcriptomic variation at the cellular level during early grain development

To investigate molecular transitions during early wheat seed development, we divided transcriptomes from 16 cell clusters into two stages, DAP 4 and DAP 8. UMAP analysis showed that all clusters were present at both stages, but their relative proportions changed markedly (Fig. 2A). Specifically, DC, ESR, TCSE, and pericarp clusters decreased, whereas AL and CSE clusters increased, indicating a developmental shift toward nutrient accumulation at DAP 8 (Fig. 2B). To characterize tissue-specific transitions, we integrated stage-specific genes identified from bulk RNA-seq and chromatin accessibility datasets with single-cell data (Zhao *et al*., 2024). Genes with high expression or accessibility showed a sequential enrichment pattern, progressing from pericarp, SC, AL, and CF at early stages to DC and ultimately CSE and TCSE at later stages (Fig. S2A and S2B). Consistently, DAP 4-upregulated genes were enriched in pericarp, SC, ESR, and DC clusters, whereas DAP 8-upregulated genes were primarily enriched in AL, TCSE, and CSE clusters, in agreement with observed shifts in cell-type composition (Fig. 2C).

**Fig. 2.**
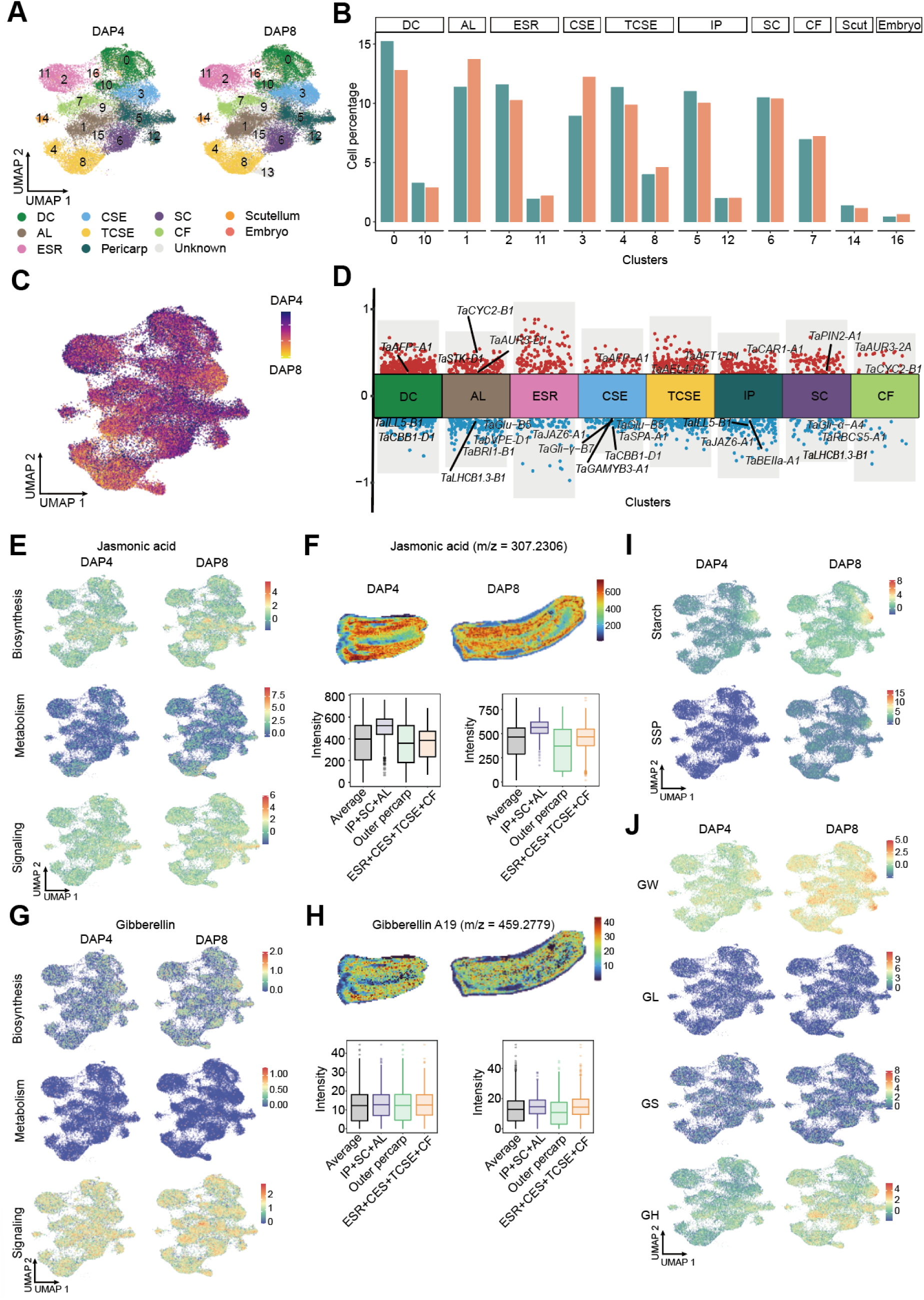
Cell-type-resolved developmental dynamics of hormone signaling and nutrient accumulation gene during early wheat grain development. A. UMAP visualization of cell types at two time points (DAP 4 and DAP 8), with cells colored by cluster identity. B. Bar charts showing the proportions of cell of in each cell cluster at DAP 4 and DAP 8. C. Developmental pseudotime trajectories for each cell cluster mapped onto the same UMAP projection shown in (A). Differentiation scores were the enrichment levels of stage-specific marker genes identified between DAP 4 and DAP 8. D. Differential gene-expression analysis showing the highly expressed genes across 7 cell-types at DAP 4 and DAP 8. DAP 4. Genes with higher expression at DAP 4 are shown in red, and those with higher expression at DAP 8 are shown in blue. E. UMAP plot shown the expression enrichment levels of jasmonic acid related biosynthetic, metabolic and signaling genes. Color-coded based on the expression levels of the selected gene sets. F. The MALDI-MS images of the jasmonic acid in Fielder grain at DAP 4 and DAP 8. The boxplot shown the quantified signal intensities across defined seed regions. G. UMAP plot shown the expression enrichment levels of gibberellin related biosynthetic, metabolic and signaling genes. Color-coded based on the expression levels of the selected gene sets. H. The MALDI-MS images of the gibberellin A19 in Fielder grain at DAP 4 and DAP 8. The boxplot shown the quantified signal intensities across defined seed regions. I. UMAP plot shown the expression enrichment levels of starch biosynthesis and seed storage protein genes. Color-coded based on the expression levels of the selected gene sets. J. UMAP plot shown the expression enrichment levels of genes associated with grain weight, grain length, grain size and grain hardness. Color-coded based on the expression levels of the selected gene sets.

Then, we performed differential expression analysis for each cell cluster, the temporal expression patterns of these genes were highly consistent with cell-type identities. For example, at DAP 4, the auxin efflux transporter *TaPIN2-A1* was highly expressed in the SC cluster; the negative regulator of seed dormancy *TaAFP-A1* showed elevated expression in the CSE cluster; and cell cycle-related genes such as *CYC2* were specifically expressed in the AL and CF clusters. By contrast, at DAP 8, genes related to starch biosynthesis and seed storage proteins were highly expressed in the AL and CSE clusters, whereas photosynthesis-related genes were specifically enriched in the SC cluster (Fig. 2D and Supplemental Data Set S5). In addition, genes involved in hormones signaling pathways exhibited pronounced cell-type-specific expression patterns. For instance, the gibberellin signaling gene *TaAEL4-D1* was specifically highly expressed in TCSE cells at DAP 4; the jasmonic acid signaling gene *TaJAZ6-A1* showed specific expression in IP cells at DAP 8; the auxin metabolism gene *TaAUR3-D1* was enriched in CF and AL clusters at DAP 4; and the brassinosteroid biosynthesis gene *TaCBB1-D1* and signaling gene *TaBRI1-B1* were specifically highly expressed in the CSE and AL clusters, respectively, at DAP 4. The Abscisic acid receptor *TaCAR1-A1* showed specific expression in IP cells at DAP 4.

Given the critical role of phytohormone distribution in regulating tissue development, we generated expression enrichment maps for genes involved in the biosynthesis, metabolism, and signaling of jasmonic acid (JA), gibberellin (GA), auxin, abscisic acid (ABA), and brassinosteroids (BRs) at DAP 4 and DAP 8 (Supplemental Data Set S6) (Feng et al., 2024). In parallel, spatial metabolite maps of JA, GA, and auxin were constructed, and signal intensities were quantitatively assessed across defined seed regions (Fig. S2C). JA biosynthetic genes were highly expressed in the IP, SC, and AL, whereas JA metabolic genes were predominantly expressed in TCSE cells. Genes involved in JA signaling displayed expression patterns similar to those of JA biosynthetic genes, with particularly strong expression in the IP (Fig. 2E). Consistently, JA metabolite spatial maps showed stronger signal intensities in the IP + SC + AL regions (Fig. 2F). In contrast, GA biosynthetic genes exhibited their highest expression levels in CSE cells at DAP 8, while GA metabolic genes showed no clear enrichment. GA signaling genes were most strongly expressed in the AL (Fig. 2G). Correspondingly, GA metabolite spatial maps revealed moderately elevated signal intensities in the ESR, CSE, TCSE and CF regions at DAP 8 (Fig. 2H). Auxin-related genes did not exhibit pronounced tissue-specific expression patterns (Figs. S2D and S2E). In addition, ABA biosynthetic genes were highly expressed in the SC, while ABA signaling genes were enriched in the IP. BR biosynthetic genes showed elevated expression in the AL at DAP 8, whereas BR signaling genes exhibited broadly increased expression at this stage (Fig. S2F). Collectively, these results demonstrate pronounced tissue-specific hormone signaling during early endosperm development, suggesting that spatially restricted hormonal regulation contributes to the development and differentiation of distinct endosperm tissues.

Furthermore, to gain insight into the formation of wheat grain yield and processing quality, we examined the enrichment of genes associated with these traits (Zhao *et al*., 2024). Genes involved in starch biosynthesis and seed storage protein (SSP) accumulation displayed markedly higher expression at DAP 8 and were strongly enriched in CSE cells (Fig. 2I). Representative examples include the *1,4-α-glucan-branching enzyme TaBEIIa-A1* and the low-molecular-weight glutenin gene *TaGlu1-B5* (Figs. S2F and S2G). In addition, genes regulating grain weight were also highly expressed in CSE cells at DAP 8, whereas genes associated with grain hardness showed higher enrichment in AL cells at this stage. By contrast, genes controlling grain length and width did not exhibit significant enrichment, suggesting that these traits may not be primarily regulated during early endosperm development (Fig. 2J). This analysis reveals dynamic, stage-specific transcriptional reprogramming across seed tissues, indicating a developmental transition from growth-related processes at DAP 4 to coordinated nutrient accumulation at DAP 8.

### Pseudotime reconstruction reveals distinct cell fate trajectories during wheat seed development

Seed development involves extensive cellular differentiation. To delineate the differentiation trajectories of wheat seed cells, we reconstructed pseudotime trajectories based on single-nucleus transcriptomic data. This analysis identified six major cell clusters and three principal branches (Fig. 3A). Most DC cells, CSE cells, and embryonic cells were located in branch cluster 7, exhibiting relatively low differentiation states. In contrast, two highly differentiated branches corresponded to distinct cell fates. One branch (cell fate 1), primarily comprising ESR and TCSE cells, was located in branch cluster 4 and represented differentiated peripheral endosperm cell lineages. The other branch (cell fate 2), mainly consisting of inner pericarp, seed coat, and endosperm cavity cells, was distributed across branch clusters 1 and 3 (Fig. 3A-C and Fig. S3A). To identify key regulatory factors underlying these two highly differentiated trajectories, we examined branch-specific gene expression dynamics along pseudotime and identified genes significantly enriched in each branch (Fig. 3D). GO enrichment analysis revealed that pathways related to protein processing, cell wall organization, cell fate determination, and brassinosteroid signaling were significantly enriched in cell fate 1, indicating pronounced cell wall thickening and functional specialization in these cells (Fig. 3E). In contrast, cell fate 2 was enriched for pathways associated with photosynthesis and nutrient transport, suggesting a role in nutrient synthesis and translocation toward endosperm tissues (Fig. 3E).

**Fig. 3.**
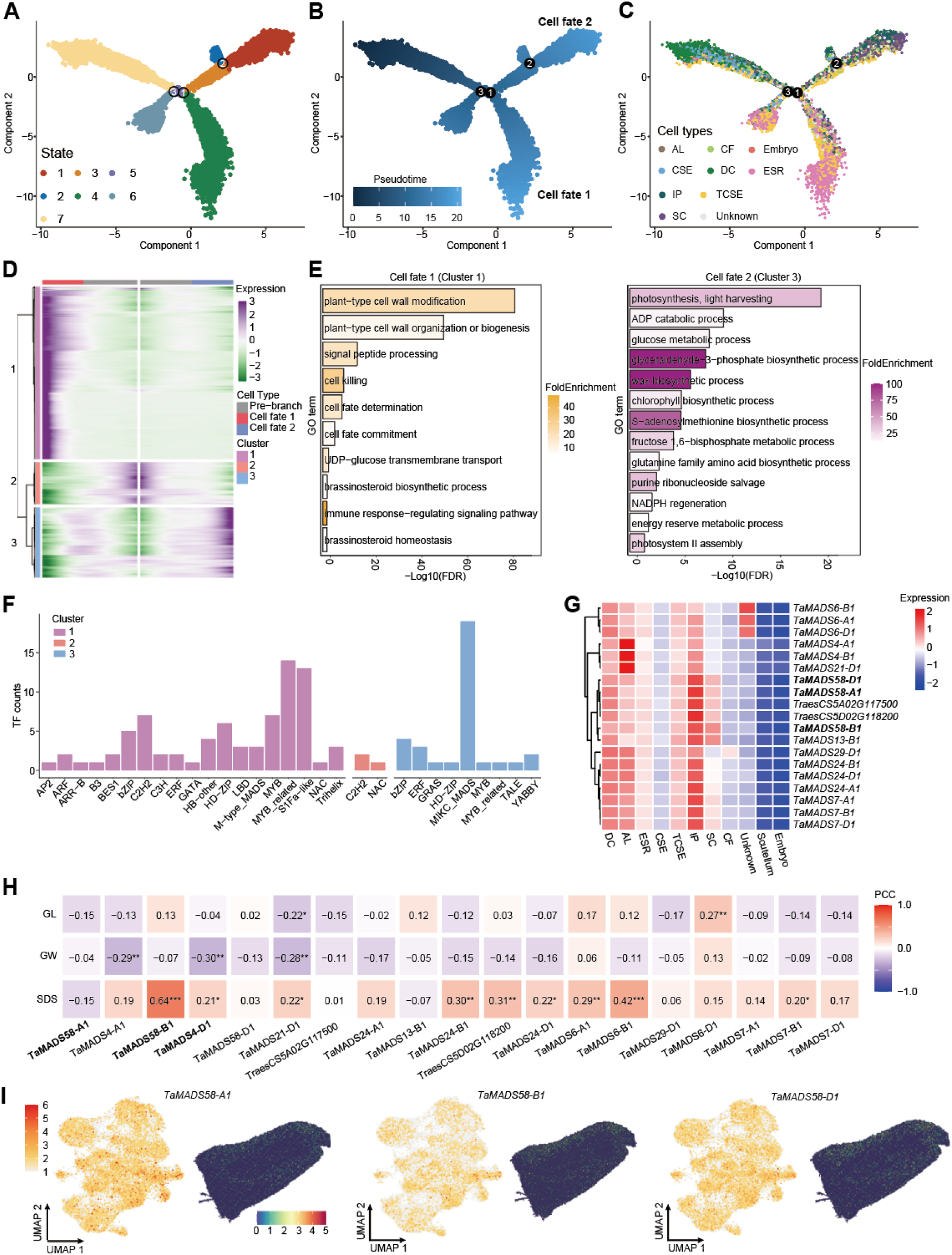
Developmental trajectories of wheat early endosperm and branch-specific gene. A-C. Visualization of the development trajectory of seed cells colored by branch (A), development stages (B) and cell types (C). D. Expression dynamics of the most highly variable (*P*-value < 0.001) genes along pseudotime between branch 1 and branch 2. E. GO enrichment for genes in (D) across cell fate 1 and cell fate 2. False discovery rate (FDR) by Benjamini-Hochberg procedure. F. The TF number in each family of genes in (D) across cell fate 1 and cell fate 2. G. The expression of MADS TF genes in Pseudo-bulk RNA-seq. H. Pearson correlation coefficients between the expression levels of MADS transcription factors (TFs) and grain-related traits, including grain length (GL), grain width (GW), thousand-kernel weight (TKW), and SDS sedimentation value (SDS). I. UMAP plot show the expression levels of *TaMADS58-A1/B1/D1*, and spatial expression patterns of *TaMADS58-A1/B1/D1*. Color-coded by expression level.

Given the central regulatory roles of transcription factors (TFs) in developmental processes, we systematically analyzed the composition and abundance of TFs among branch-specific genes (Fig. 3F and Supplemental Data Set S7). Notably, MADS-box TFs were significantly enriched in branch 2, implying their involvement in establishing nutrient transport during seed development. Pseudo-bulk RNA-seq analysis derived from single-cell data further classified these MADS TFs into two groups with preferential expression in the aleurone layer and inner pericarp cells, respectively, consistent with the inferred differentiation trajectories and supporting their cell-type-specific regulatory roles (Fig. 3G). To further investigate the biological significance of these MADS TFs, we examined the association between their expression levels and key grain traits using previously published population-scale grain transcriptome datasets (Zhao *et al*., 2024). (Fig. 3H). Among them, the expression of *TaMADS58-B1* showed a significant positive correlation with the SDS sedimentation value, a key indicator of wheat processing quality (Fig. 3H and S3B), suggesting a potential role in grain quality regulation. However, population-scale transcriptome analyses did not reveal a significant association between these homoeologs and SDS sedimentation value (Fig. S3B and S3C), likely because *TaMADS58-A1* and *TaMADS58-D1* lack haplotypes with substantial expression variation in the analyzed population (Fig. S3B). Both single-cell UMAP visualization and spatial transcriptomic analyses revealed high expression of *TaMADS58s* in pericarp tissues, with *TaMADS58-A1* exhibiting the highest expression level. This indicates that it may influence grain quality through regulation of pericarp development (Fig. 3I).

Collectively, these findings elucidate the differentiation trajectories and cell fate decisions during wheat seed development and highlight a critical role for MADS-box transcription factors, particularly *TaMADS58s* in regulating nutrient transport pathways that ultimately impact wheat grain yield and quality.

### *TaMADS58* regulates pericarp development while negatively modulating grain size and enhancing wheat end-use quality

To validate the role of *TaMADS58* in wheat seed development, we generated both *TaMADS58-A1* overexpression (OE) lines in the “Fielder” background and *TaMADS58-B1* loss-of-function mutants (*tamads58-b1*) in the “KN9204” background (Fig S4A). Transverse sections of developing seeds revealed a pronounced increase in pericarp thickness in *TaMADS58-A1*-OE lines at both DAP 4 and DAP 8 (Fig. 4A). In contrast, the pericarp was significantly thinner in *tamads58-b1* (Fig. 4B). Given that the wheat pericarp is progressively degraded during grain development, these results suggest that TaMADS58 may regulate pericarp thickness by repressing pericarp degradation. Phenotypic analysis of mature grains showed that both grain length and grain width were significantly reduced in *TaMADS58-A1*-OE lines, whereas they were significantly increased in *tamads58-b1* (Fig. 4C, Fig. S4B and S4C). In addition, quality-related traits were altered in the *TaMADS58-A1*-OE, including increased total protein and gluten content accompanied by a reduction in total starch content (Fig. 4D). These findings indicate that *TaMADS58* positively regulates grain quality traits but negatively affects starch accumulation and yield-related traits.

**Fig. 4.**
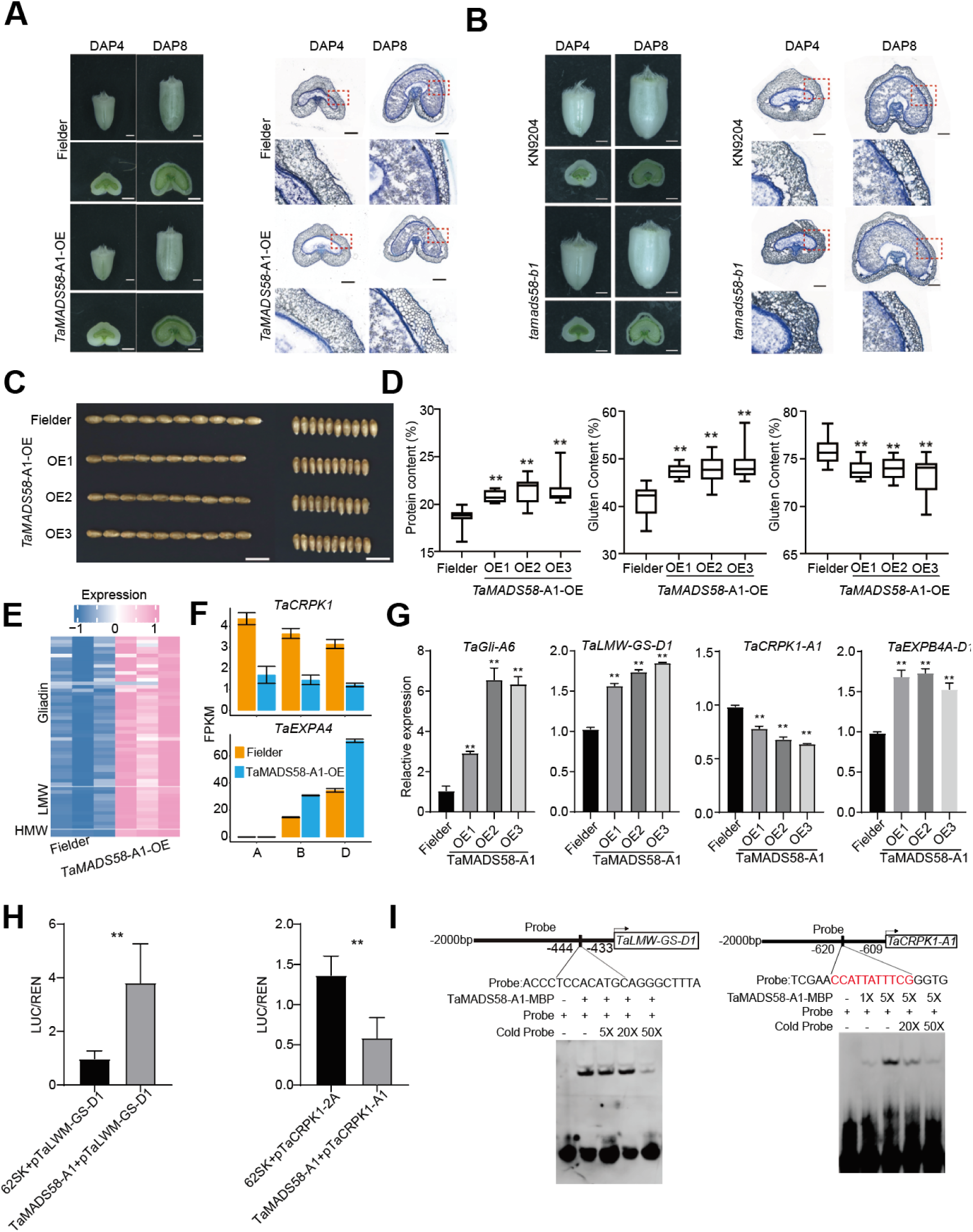
*TaMADS58* regulates regulates the thickness of the pericarp of wheat grains and seed storage proteins. A. Developing grains phenotypes of Fielder (WT) and *TaMADS58-A1*-OE (left). A representative section of the middle portion of a developing grain of Fielder (WT) and *TaMADS58-A1-OE* wheat (right). Bars=200 μm. B. Developing grains phenotypes of KN9204 (WT) and *tamads58-b1* (left). A representative section of the middle portion of a developing grain of KN9204 (WT) and *tamads58-b1* wheat (right). Bars=200 μm. C. Grain traits of the *TaMADS58-A1*-OE. The grains of different lines exhibited significant differences in length and width. N = 10 grains, Bars = 1 cm. D. Quantification of grain agronomic traits related traits between the WT plants and *TaMADS58-A1-OE* lines. Student’s t test was used to determine the difference significance between *TaMADS58-A1*-OE and WT. *, *P* ≤ 0.05, **, *P* ≤ 0.01, Data represent mean ± SD (n = 15 biological replicates). E. Heatmap showing the expression levels of gliadin, glutenin genes in Fielder and Ta*MADS58-A1*-OE line. F. Expression levels of *TaCRPK1-A1/B1/D1* and *TaEXPA4-A1/B1/D1* between *TaMADS58-A1*-OE and Fielder. Data are presented as means ± SD. n = 3. G. Expression analysis of *TaMADS58-A1*-OE and its downstream target genes in WT and OE1 lines. *TaACTIN* was used as the internal control. Data are presented as means ± SD. *P*-values were determined by Student’s t-test. n = 3. H. Transient expression assays in N. benthamiana leaves show that TaMADS58-A1 activates *TaLMW-GS-D1* suppresses *TaCDPK1-A1*. Relative luciferase activity is expressed as the LUC/REN ratio. I. Electrophoretic mobility shift assays (EMSAs) show that *TaMADS58-A1* specifically recognizes the CArG element in the promoters of *TaLWM-GS-D1* and *TaCDPK1-A1*. “Probe” indicates biotin-labeled DNA, “competitor” denotes unlabeled DNA, and “mCompetitor” refers to a mutated motif without biotin labeling.

Subcellular localization assays demonstrated that *TaMADS58* is localized in the nucleus, consistent with its putative role as a transcription factor (Fig. S4D). To further investigate its regulatory function, we performed RNA-seq analysis on DAP 8 seeds from *TaMADS58-A1*-OE and wild-type ‘Fielder’ (Fig. S4E). A large number of *seed storage protein* (*SSP*) genes were upregulated in the OE lines, whereas only a single starch biosynthesis gene, *TaAGPS1*, was downregulated (Fig. 4E and Supplemental Data Set S8), indicating a predominant positive regulatory role of *TaMADS58* in SSP biosynthesis with minimal impact on starch biosynthetic pathways. GO enrichment analysis further revealed that genes associated with cell wall biosynthesis and carbohydrate metabolism were significantly upregulated, while stress-response genes were downregulated in the *TaMADS58-A1*-OE (Fig. S4F). Previous studies have demonstrated that *TaEXPA4*-*D1* positively regulates pericarp development, whereas *Zm00001d002283* (the wheat homolog *TaCRPK1*) functions as a negative regulator of pericarp development (Calderini et al., 2021; Gong et al., 2024). In *TaMADS58-A1*-OE, *TaEXPA4-B1/D1* expression was significantly increased (*TaEXPA4-A1* was not expressed), while *TaCRPK1-A1/B1/D1* expression was markedly decreased, suggesting that TaMADS58 may modulate pericarp development through coordinated regulation of these genes. To validate these expression changes, we selected several representative genes for quantitative PCR (qPCR) analysis, which confirmed the upregulation of several SSP genes, including TaGli-6 and *TaLMW-GS-D1*, as well as *TaEXPA4-D1*, together with the downregulation of *TaCRPK1-A1* (Fig. 4G). Furthermore, luciferase reporter assays, electrophoretic mobility shift assays (EMSA), and yeast one-hybrid assays demonstrated that *TaMADS58-B1* directly regulates the expression of *TaLMW-GS-D1* and *TaCRPK1-A1* (Fig. 4H, 4I, and Fig. S4G).

In summary, we demonstrate that *TaMADS58* directly activates the expression of SSP encoding genes while simultaneously delaying pericarp degradation by repressing *TaCRPK1-A1*.

### Single-cell co-expression networks reveal cell type-specific regulatory modules

The translation of gene expression into biological function relies on hierarchical regulatory networks, in which genes involved in similar biological processes often exhibit coordinated expression patterns. Previous co-expression network analyses based on bulk RNA-seq data have successfully identified functionally related regulatory modules; however, their resolution is limited by cell-type heterogeneity, restricting the granularity of inferred modules. To overcome this limitation, we performed a single-cell-level co-expression network analysis using hdWGCNA, applying an iterative network construction strategy to the snRNA-seq dataset (Morabito et al., 2023). Across 11 distinct cell clusters, we identified 11 co-expression modules with pronounced cell-type specificity (Fig. 5A). Genes within each module exhibited clear tissue-enriched expression patterns (Fig. 5A). For example, the red module was enriched in ESR cells and contained the grain-weight-associated gene *Dolichyl-diphosphooligosaccharide-protein glycosyltransferase subunit* gene *STT3* (*TaSTT3a-A1*). The greenyellow module was specifically enriched in CF cells and included the *vacuolar processing enzyme* gene *TaVPE1-A1*. The pink and purple modules were preferentially expressed in TCSE cells and harbored genes such as the *lipid-transfer protein* gene *TaLTPL36-A1* and the *plasma membrane ATPase* gene *TaAHA7-B1*. The magenta, brown, and green modules were enriched in DC cells and contained cell cycle-and chromatin-related genes, including *TaCycB1;1-B1* and *Histone H2A* (*TaH2A-A1*). The blue module was enriched in SC cells and included *Glucose-1-phosphate adenylyltransferase* gene (*TaAGPL2-A1*), whereas the black module was enriched in IP cells and contained *Fructose-bisphosphate aldolase* gene (*TaFBA3-A1*) (Fig. 5B, 5C, Fig. S5A and Supplemental Data Set S9).

**Fig. 5.**
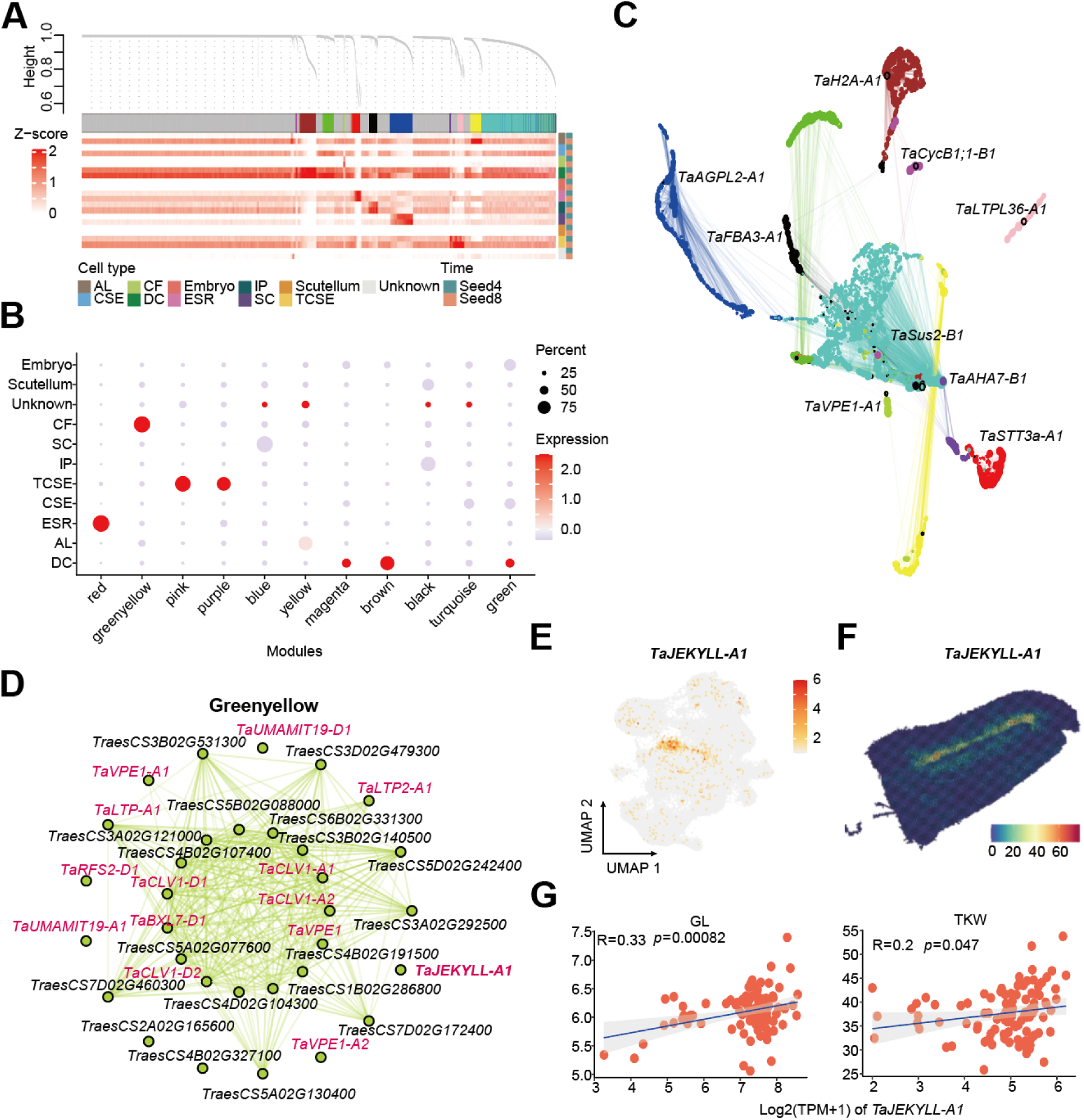
Single-cell co-expression networks and identification of the cell-type-specific regulator *TaJEKYLL-A1*. A. Dendrogram showing the hierarchical clustering of isoforms into co-expression modules based on the topological overlap matrix (TOM). The heatmap shown the expression of gene from each module in single-cell data simulated bulk RNA-seq. B. Single-cell co-expression networks. Each node represents a single gene, and edges represent co-expression links between genes and module hub genes. Nodes are coloured by co-expression module assignment. The top one hub genes per module are labelled. C. Dot plot presenting the average expression of module-specific hub genes in different clusters, calculated by gene scoring for the top 20 hub genes ranked by kME in each module. D. Network plot displays the hub genes in the greenyellow module. The known genes are highlight. E. UMAP plot show the expression levels of *TaJEKYLL-A1*. F. Spatial expression patterns of *TaJEKYLL-A1*.

Among these modules, the greenyellow module displayed particularly strong specificity for CF cells. The CF contains transfer cell types such as the nucellar projection, which are essential for transporting maternal carbon and nitrogen resources into the developing grain and therefore play a critical role during endosperm development. To validate this observation, we examined the expression patterns of genes involved in carbon and nitrogen transport. The results showed that sugar transporters of the SWEET family and amino acid transporters of the UMAMIT family were specifically expressed in CF cells (Fig. S5B and Supplemental Data Set S10) (Guo et al., 2025), further highlighting the importance of CF cells in nutrient transport. To identify potential key regulators of nucellar projection and CF cell development, we focused on hub genes within the greenyellow module. This module was enriched for multiple vacuolar processing enzymes gene (*VPEs*), the amino acid transporter gene *TaUMAMIT19-D1*, the lipid-transfer protein gene *TaLTP2-A1*, several leucine-rich repeat receptor-like kinases genes, and *TaJEKYLL-A1* (Fig. 5D). Notably, *JEKYLL* has been reported in barley to regulate nucellar projection development (Radchuk et al., 2019). Consistently, *TaJEKYLL-A1* exhibited CF-specific expression patterns in both snRNA-seq and spatial transcriptomics datasets (Fig. 5E and 5F), suggesting a conserved role in wheat. Furthermore, population-level transcriptome analyses revealed that the expression level of *TaJEKYLL-A1* was significantly correlated with grain length and thousand-kernel weight (Fig. 5G), implying its potential contribution to wheat yield regulation.

These results demonstrate that single-cell-resolved co-expression network analysis uncovers cell-type-specific regulatory modules in wheat grains and identifies *TaJEKYLL-A1* as a promising candidate regulator of nucellar projection development and nutrient transport.

### *TaJEKYLL* regulates the development of nucellar projection, affecting nutrient transport from maternal tissue to seeds

To further elucidate the biological function of *TaJEKYLL*, we first confirmed its cell-type specificity by in situ hybridization, which demonstrated that *TaJEKYLL* is specifically expressed in the nucellar projection cells of the endosperm cavity (Fig. 6A). We then generated *TaJEKYLL-A1/B1* double-knockout lines (*jekyll-abD*) and a *TaJEKYLL-A1* single-mutant line (*jekyll-a1*) (Fig. S6A). Mature grains of both *jekyll-abD* and *jekyll-a1* were significantly smaller than those of the wild type, suggesting impaired nutrient transport following *TaJEKYLL* disruption (Fig. 6B and S6B). Transverse sections of developing seeds further revealed a pronounced reduction in the size of the nucellar projection in both *jekyll-abD* and *jekyll-a1* (Fig. 6C and S6C). Quantitative analyses confirmed significant decreases in both the width and area of the nucellar projection in *jekyll-abD* and *jekyll-a1* (Fig. 6D and S6D), indicating that *TaJEKYLL* is required for proper nucellar projection development. Consistently, the expression levels of the sucrose transporters *TaSWEET11* and *TaSUT1* were markedly reduced in *jekyll-abD*, implying compromised sucrose transport into the developing grain (Fig. 6E). In addition, several starch biosynthesis-related genes, including *TaAGPL1*, *TaSUS1*, *TaSUS7*, and *TaAGPS1a*, were significantly downregulated in *jekyll-abD* (Fig. 6E and S6E), suggesting that defective nucellar projection development ultimately impairs starch accumulation and leads to reduced grain size. Previous studies have shown that *TaMADS29* regulates nucellar projection development by controlling programmed cell death (Liu *et al*., 2023). We therefore investigated whether *TaMADS29-A1* acts upstream of *TaJEKYLL-A1* to regulate its expression (Fig. 6F and 6G). Yeast one-hybrid and luciferase reporter assays demonstrated that TaMADS29-A1 activates *TaJEKYLL-A1* transcription, while EMSA confirmed that TaMADS29-A1 directly binds to the *TaJEKYLL-A1* promoter in vitro (Fig. 6H).

**Fig. 6.**
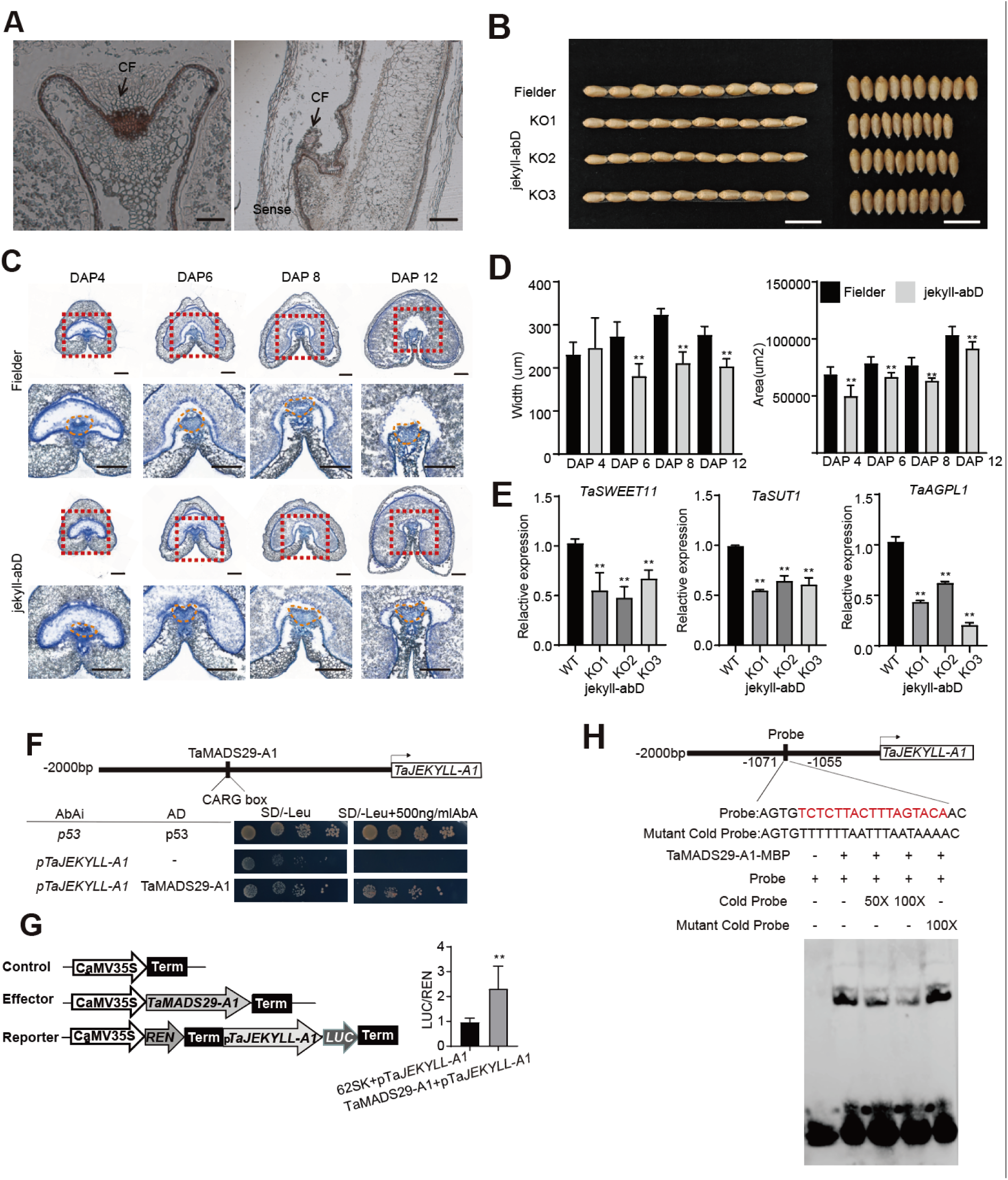
*TaJEKYLL* negatively regulates wheat nutrient transfer. A. Expression pattern of *TaJEKYLLs* in grain at DAP 12 as detected by RNA in situ hybridization. CF, cavity fuild. Scale bars, 100 μm. B. Grain traits of the *tajekyll-abD.* The grains of different lines exhibited significant differences in length and width. N = 10 grains, Bars = 1 cm. C. Transverse section of Developing grains revealed the nucellar projection degradation in the *tajekyll-abD*. Red boxes in the top panel were magnified and represented in the lower panel. Bars=200 μm. D. Statistic comparison of nucellar projection width and area between Fielder and *tajekyll-abD*. n = 3. Significance was determined using Student’s t-test, **P < 0.01. E. Expression analysis of *tajekyll-abD* and its downstream target genes in WT and KO1 lines. *TaACTIN* was used as the internal control. Data are presented as means ± SD. *P*-values were determined by Student’s t-test. n = 3. F. Y1H assay of TaMADS29-A1 binds to *TaJEKYLL-A1* promoter. G. Transient expression assays in N. benthamiana leaves show that TaMADS29-A1 suppresses *TaJEKYLL-A1*. Relative luciferase activity is expressed as the LUC/REN ratio. H. Electrophoretic mobility shift assays (EMSAs) show that *TaMADS29-A1* specifically recognizes the CArG element in the promoters of *TaJEKYLL-A1*. “Probe” indicates biotin-labeled DNA, “competitor” denotes unlabeled DNA, and “mCompetitor” refers to a mutated motif without biotin labeling.

Overall, *TaJEKYLL* acts as a positive regulator of nucellar projection development, thereby promoting efficient nutrient transport and contributing to increased grain size.

### Natural variation of *TaMADS58-B1* and *TaJEKYLL-A1* and their contributions to global wheat yield and end-use quality

To systematically characterize natural sequence variation, expression divergence, and their associations with grain-related traits, we analyzed coding-region polymorphisms of *TaMADS58-B1* and *TaJEKYLL-A1* using previously published wheat population transcriptome datasets (Zhao *et al*., 2024). Three single-nucleotide polymorphisms (SNPs) were identified in the coding region of *TaMADS58-B1*, based on which wheat accessions were classified into two haplotypes (Figs. 7A and 7B). Hap1 exhibited significantly higher expression levels (Fig. 7C) and was associated with end-use quality traits, including MPT_MRW (peak time × post-peak bandwidth), MPT_MTW (peak time × tail bandwidth), and SDS sedimentation value, compared with Hap2. In contrast, no significant differences were observed between the two haplotypes in grain length or thousand-kernel weight (Fig. 7D). These results indicate that Hap1 of *TaMADS58-B1* enhances wheat processing quality without compromising yield. In the coding region of *TaJEKYLL-A1*, two SNPs were identified, allowing classification of the germplasm into two haplotypes (Figs. 7E and 7F). Hap1 showed a modestly higher expression level (Fig. 7G). Although no significant differences were detected between haplotypes in end-use quality traits (MPT_MRW, MPT_MTW, and SDS sedimentation value), accessions carrying Hap1 displayed significantly increased grain length and thousand-kernel weight (Fig. 7H). This suggests that Hap1 of *TaJEKYLL-A1* primarily contributes to yield improvement without negatively affecting processing quality.

**Fig. 7.**
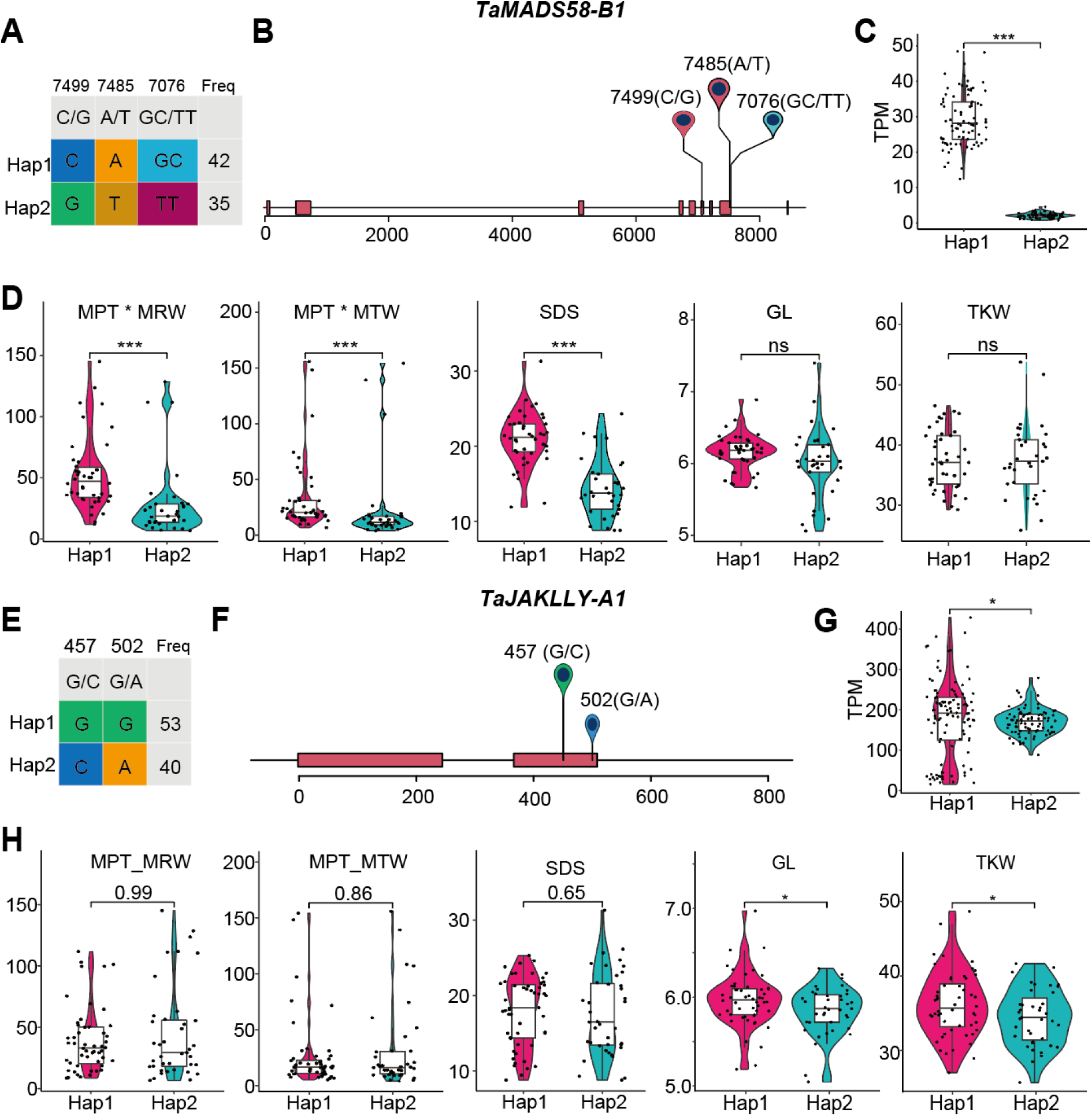
Haplotype analysis of *TaMADS58-B1* and *TaJEKYLL-A1* and their association with grain traits. A. The SNPs and haplotypes of *TaMADS58-B1*. B. Schematic showing the polymorphism for each haplotype of *TaMADS58-B1*. The coordinate is related to the transcription start site (TSS) C. Violin plot indicating the comparison of expression levels among wheat accession with different haplotypes of *TaMADS58-B1*. Statistical significance was assessed by Student’s *t*-test. D. Violin plot indicating the comparison of end-use quality-related traits MPT_MRW (peak time × post-peak bandwidth), MPT_MTW (peak time × tail bandwidth), and SDS sedimentation value and yield-related traits grain length and thousand-kernel weight among wheat accession with three haplotypes of *TaMADS58-B1*. Statistical significance was assessed by Student’s *t*-test. E. The SNPs and haplotypes of *TaJEKYLL-A1*. F. Schematic showing the polymorphism for each haplotype of *TaJEKYLL-A1*. The coordinate is related to the transcription start site (TSS) G. Violin plot indicating the comparison of expression levels among wheat accession with different haplotypes of *TaJEKYLL-A1*. Statistical significance was assessed by Student’s *t*-test. H. Violin plot indicating the comparison of end-use quality-related traits MPT_MRW (peak time × post-peak bandwidth), MPT_MTW (peak time × tail bandwidth), and SDS sedimentation value and yield-related traits grain length and thousand-kernel weight among wheat accession with three haplotypes of *TaJEKYLL-A1*. Statistical significance was assessed by Student’s *t*-test.

To further investigate genetic variation in noncoding regions, we performed polymorphism analyses of the coding sequence, 3-kb upstream promoter region, and 3-kb downstream region of *TaMADS58-B1* and *TaJEKYLL-A1* using accessions from the Watkins wheat collection (Cheng et al., 2024). For *TaMADS58-B1*, one SNP was detected in the promoter region, no SNPs were identified in the coding region, and two SNPs were found in the 3′ downstream region, revealing a variation pattern distinct from that observed in the population transcriptome dataset (Fig. S7A). Based on these variants, accessions were grouped into three haplotypes: 327 accessions carrying Hap1, 271 carrying Hap2, and 60 carrying Hap3 (Fig. S7B). Hap1 predominated in Asia, whereas Hap2 was more frequent in Europe (Fig. S7C). In *TaJEKYLL-A1*, 14 SNPs were identified in the promoter region, two SNPs in the coding region (consistent with transcriptome-based analyses), and 17 SNPs in the 3′ downstream region (Fig. S7D). These variants defined three major haplotypes: Hap1 (352 accessions), Hap2 (153 accessions), and Hap3 (149 accessions) (Fig. S7E). Hap1 predominated in Asia and Western Europe, whereas Hap2 was most frequent in the Mediterranean region (Fig. S7F).

Collectively, these results suggest that regional ecological conditions may have driven the selection and dissemination of specific alleles of *TaMADS58-B1* and *TaJEKYLL-A1*, thereby facilitating coordinated adaptation and improvement of wheat yield and quality across diverse environments.

## Discussion

Wheat grain development is characterized by pronounced cellular specialization, with distinct cell types performing divergent functions within a tightly organized tissue architecture. However, transcriptional programs derived from bulk RNA-seq inherently represent population averages, obscuring functionally relevant cell-type heterogeneity within highly differentiated organs. Single-cell /single-nucleus transcriptomic approaches overcome this limitation by resolving mixed cell populations into discrete cellular states, enabling systematic annotation of cell identities, quantification of transcriptional heterogeneity, and reconstruction of developmental trajectories through pseudotemporal analysis (Lee et al., 2025). Despite these advances, single-cell datasets for wheat grains remain limited, and cellular identities are frequently disconnected from their native spatial context.

Integrating single-cell transcriptomics with spatial transcriptomic frameworks addresses this gap by anchoring cell types within anatomical coordinates, revealing cellular identity, spatial organization, and function during cereal grain development. This combined strategy has already been rapidly adopted across crops (Wang et al., 2024; Zhang et al., 2025c; Zhu et al., 2025).

By integrating Stereo-seq and single-cell RNA-seq datasets from two key stages of wheat seed development, we generated a spatially and temporally resolved transcriptomic atlas approaching single-cell resolution. This integrative framework substantially expanded the detectable transcriptome beyond bulk RNA-seq, enabling the resolution of cell-type-specific expression and the discovery of distinct subgenome-biased gene expression patterns. (Fig. 1E-F). Polyploidization is frequently associated with asymmetric gene expression among subgenomes and their unequal contributions to biological processes (Feldman et al., 2012), and previous ATAC-seq and RNA-seq studies have revealed pronounced stage-dependent subgenome bias in transcription factor activity, with A-subgenome dominance during early endosperm development and increasing contributions from the B and D subgenomes at later grain-filling stages (Pei et al., 2023). These observations underscore the importance of resolving gene expression at both cellular and spatial resolution to elucidate how individual subgenomes are differentially deployed during seed development in polyploid crops. Although spatial transcriptomics enables the analysis of subgenome expression within intact tissue architecture, its relatively limited transcript capture depth has resulted in the identification of only a small number of subgenome-biased candidates in previous studies (Li *et al*., 2025; Millsteed et al., 2025). Single-cell RNA sequencing offers higher cellular resolution and greater transcriptome coverage, allowing robust detection of lowly expressed genes and subtle transcriptional differences among closely related cell types. Leveraging snRNA-seq, we identified 1,147 genes exhibiting clear subgenome-biased expression across distinct cell clusters, indicating that subgenome expression asymmetry is tightly linked to cellular heterogeneity.

Our analysis revealed pronounced cell-type-specific expression patterns for genes involved in hormone signaling pathways (Fig. 2 E-F). Using spatial metabolomic imaging, we further visualized tissue-level distributions of JA, GA, and auxin in developing wheat grains (Fig. 2 G-H, Figs. 2 E-F). Previous studies have leveraged single-cell and spatial transcriptomic approaches to map cell-type-specific expression of hormone biosynthesis and signaling genes during rice seed germination and wheat spike development, revealing dynamic and cell-type-restricted regulatory programs (Yao *et al*., 2024; Zhang *et al*., 2025c). However, direct high-resolution visualization of hormone molecules within developing seeds has remained largely unexplored. By integrating single-cell transcriptomics with spatial metabolomic imaging, we provide the first tissue-level maps of JA, GA, and auxin distributions in developing wheat grains. This approach demonstrates that, while hormone biosynthetic and signaling genes show cell-type-specific expression, the actual hormone molecules form spatially restricted hotspots or gradients (Fig. 2 E-H). Collectively, these results underscore the value of combining gene expression profiling with in situ hormone visualization to capture the full complexity of developmental signaling. Meanwhile, our data were further integrated through bulk RNA-seq, ATAC-seq, and snRNA-seq analyses (Fig. S2 A-B) (Zhao *et al*., 2024; Zhao *et al*., 2023), enabling the construction of cell developmental atlases that provide valuable insights into the spatial regulation of development and the coordination of cellular differentiation programs.

Pseudotime analysis revealed an enrichment of MADS-box transcription factors along developmental trajectories associated with nutrient transport, highlighting their coordinated involvement in seed maturation programs. Among these, TaMADS58 emerged as a central regulatory node whose expression was positively correlated with SDS sedimentation value and grain protein content (Fig. 3I). TaMADS58 promoted the expression of albumin- and glutenin-encoding genes (Fig. 4D), directly linking its transcriptional activity to storage protein biosynthesis. Notably, TaMADS58 also modulated the expression of key regulators of pericarp degradation, including TaLWM and TaCRPK1, establishing a mechanistic connection between maternal tissue remodeling and filial nutrient accumulation. These results therefore extend pericarp degradation from a passive developmental outcome to an actively regulated process embedded within a transcriptional network that integrates maternal tissue dynamics with filial storage programs. This consist with previous studies demonstrating that maternal tissues of cereal seeds, including the pericarp and nucellar projections, undergo tightly regulated programmed cell death in a temporally and spatially coordinated manner, facilitating nutrient remobilization and accommodating endosperm expansion (Domínguez and Cejudo, 2014; Rasheed et al., 2017; Young and Gallie, 2000).

Against this regulatory backdrop, the nucellar projection (NP) has long been recognized as a pivotal conduit for maternal-to-filial nutrient transfer (Domínguez et al., 2001; Peukert et al., 2014).Unlike maize and rice, wheat forms a unique structure groove along the ventral side of the grain, where the NP develops on the inner side and serves as the primary channel connecting maternal tissues to the endosperm (Xurun et al., 2015). Our single-cell transcriptomic analysis revealed exceptionally high and cell-type-specific expression of genes encoding sugar and amino acid transporters, as well as the monocot-specific JEKYLL protein in NP cells, with peak expression coinciding with the grain-filling phase.

Moreover, the transcriptional programs of nucellar projection (NP) cells are distinct yet complementary to those of adjacent endosperm aleurone cells. NP cells provide a critical pathway for transporting nutrients (sugars and minerals) from the mother plant, whereas aleurone cells specialize in nutrient storage and redistribution (Liang et al., 2025; Liu et al., 2025; Sheraz et al., 2021). Notably, programmed cell death (PCD) in NP cells is essential for sustaining nutrient flux (Domínguez *et al*., 2001; Yin and Xue, 2012). Consistent with this, tamads29 mutants exhibit premature PCD in both the endosperm and NP, accompanied by cell collapse and impaired maternal-to-filial nutrient transport, ultimately resulting in seed shrinkage (Liu *et al*., 2023). Our data further position JEKYLL as a downstream target of TaMADS29, modulating early maternal-filial nutrient flux and influencing grain filling, thereby affecting final grain weight. This spatial and functional dissection of NP cells deepens our mechanistic understanding of maternal-endosperm interactions and highlights NP development and activity as a potential rate-limiting determinant of grain filling efficiency and final yield. This study provides a high-resolution spatiotemporal atlas of wheat grain development, integrating single-nucleus and spatial transcriptomics with epigenomic and metabolomic profiling. By linking cell-type-specific transcriptional programs, nutrient transport pathways, and maternal-filial interactions to grain protein accumulation, these findings establish a comprehensive framework for understanding and improving cereal grain development.

## Methods

### Plant growth and tissue preparation

Seeds of winter wheat (KN9204) were germinated and subsequently transferred to soil, where they were grown at 4°C in the dark for 45 days. Then they were transferred to a greenhouse (20°C-22°C, 16-h light/8-h dark).

The EMS mutants from the tetraploid variety “KN9204” were produced by Dr. Jun Xiao’s group at the Institute of Genetics and Developmental Biology, Chinese Academy of Sciences, and were backcrossed two times before analysis.

*TaMADS58-A1*-OE T2 generation lines were grown in the experimental field of the Peking University Institute of Advanced Agricultural Sciences, Shandong, China (119°18’ E, 36°16’ N). For the field experiment, each line was planted in three replicates. Each replicate consisted of 10 rows, each 1.5 meters long, with a row-to-row spacing of 0.20 meters.

For agronomic trait measurements, grains were harvested from the spike, and 150 grains were randomly selected for analysis. A normal distribution analysis of the selected grains confirmed that grain sizes followed a normal distribution, ensuring the representation of grains from both the apical and basal regions. Grain width, grain length, and hundred-grain weight were measured using a Wanshen SC-G seed detector (Hangzhou Wanshen Detection Technology Co., Ltd.). Grain length and width were measured for more than 70 grains. Protein content and total starch content were measured using a near-infrared spectrum instrument (DA7250) with over 150 grains.

### Nucleus isolation

The nuclei isolation procedure was modified from the product information sheet of the CelLyticTM PN Isolation/Extraction Kit (Sigma). Briefly, plant tissues were chopped and mixed into 1× NIBTA buffer (1× NIB, 1 mM dithiothreitol, 1× ProtectRNATM RNase inhibitor, 1× cOmplete^TM^, ethylenediaminetetraacetic acid-free Protease Inhibitor Cocktail, and 0.3% Triton X-100) on prechilled plates, and homogenates were transferred to 15 mL tubes. The tubes were then shaken for 5 minutes on ice, after which lysates were passed through 40 μm strainers, and flow-throughs were collected into new 15 mL tubes. These were then centrifuged for 10 minutes at 1,260 × g, the supernatants were decanted, and the pellets were resuspended in 4 mL of 1× NIBTA buffer. In new 15 mL tubes containing 80% Percoll® solution (4 mL Percoll® plus 1 mL NIBTA buffer), lysates were carefully overlaid onto the Percoll® layers. Tubes were centrifuged for 30 minutes at 650 × g, after which most of the nuclei had banded at the 1× NIBTA buffer and Percoll® interface. The nuclei bands were gently collected into new 15 mL tubes, 10 mL of 1× NIBTA buffer was added, and the tubes were recentrifuged for 5 minutes at 1260 × g. Nuclei pellets were then washed twice in 1× NIBTA buffer and resuspended in PBS containing 0.04% BSA (for snRNA-Seq) to a final concentration of 2,000 nuclei/uL.

### Single-RNA sequencing

The snRNA-seq libraries were prepared using the DNBelab C Series High-throughput Single-Cell RNA Library Preparation Kit (MGI, 940-000047-00) as previously described (Liu et al., 2019). Sequencing was performed on MGI 2000, DNBSEQ T7, and DIPSEQ T1 plantforms

### Cell-type clustering and marker-gene identification

The snRNA-seq data at DAP 4 and 8 samples were aligned and counted using the Cell Ranger pipeline (version 4.0). For additional data analysis, the raw count matrix was imported into R using the Seurat (version 4.3.3) package (Hao et al., 2021). Cells with fewer than 2000 features were filtered, as were genes with expression less than 3 cells were filtered. The data was then normalized using the “SCTransform” function. We performed principal component analysis (PCA) using the “RunPCA” function and retained 30 principal components. The all datasets were integrated using Seurat. The combined seed dataset was first split into separate objects according to developmental time points using the “SplitObject”. Highly variable features for integration were identified using “SelectIntegrationFeatures”, and the datasets were subsequently prepared for SCT-based integration with “PrepSCTIntegration”. Integration anchors across time points were finally identified using the “FindIntegrationAnchors” with the normalization method set to “SCT”. Finally, the datasets were integrated using the IntegrateData function based on these anchors to generate a unified, batch-corrected expression matrix. Subsequent analyses were performed on the integrated gene-cell matrix. clustered according to resolution = 0.5, and the clustering binning results visualized using the Seurat functions RunUMAP (dim = 1:30), respectively. In order to address the impact of cell cycle heterogeneity on cell clustering, the “CellCycleScoring” function was utilized to determine the cell cycle score for each individual cell, based on the cycling orthologous genes found in rice and maize known cycling gene (Supplemental Data Set S2). The function FindAllMarkers (Wilcoxon rank-sum test, min.pct = 0.1, logfc.threshold = 0.25) was used to find all the marker genes in each cluster.

### Cellular differentiation score

Temporal differential expression for DAP 4 and 8 was analyzed using the Seurat package’s FindAllMarkers (Wilcoxon rank-sum test, min.pct = 0.1, logfc.threshold = 0.25) function. The gene signature scores (GSCs) of DAP 4 and 8 differential expression genes were calculated using UCell (version 1.3.1) (Andreatta and Carmona, 2021). The cellular differentiation score was calculated as a log2 ratio of all DAP 8/DAP 4 differential expression GSCs.

### Spatial transcriptome analysis

The spatial transcriptome data from wheat grain in DAP 4, 8 and 12 were download from spatial transcriptomic dataset (http://omicsplant.cn/WheatDB/index.html) (Li *et al*., 2025). The gene expression levels are visualized using R package ggplot2 (version 3.5.2).

### Subgenome asymmetric expression analysis

The asymmetric expression analysis of bulk RNA-seq and snRNA-seq were based on the gene triads across the homoeologs of A, B, and D subgenomes. A total of 17,311 syntenic homoeolog triads for 51,933 genes were extracted from the previous publication (Ramírez-González et al., 2018) and used for further analysis. To standardize the relative expression of each homoeolog gene across the triad, we normalized the expression level so that the sum of the expression of the three genes in each triad is 1. After the Euclidean distance of each triad was calculated, we classified the triads into seven categories as previously defined, i.e., Balance, A.dominant, B.dominant, D.dominant, A.suppressed, B.suppressed, and D.suppressed (Ramírez-González *et al*., 2018).

### GO enrichment analysis

GO enrichment was performed in GENEONTOLOGY (https://geneontology.org/), and the GO terms were visualized using R package ggplot2 (version 3.5.2).

### Pairwise comparisons of endosperm cell clusters in wheat and maize

The average expression of orthologous marker genes in each cell cluster from wheat and the published snRNA-seq datasets in maize (Yuan *et al*., 2024) were used to perform the cross-species pairwise cluster comparisons as described. The orthologous genes among wheat and maize, were downloaded from the Triticeae-Gene Tribe (http://wheat.cau.edu.cn/TGT/index.html) (Supplemental Data Set S3) (Chen et al., 2020). The number of overlapped orthologous genes for each cluster was visualized using R package ggsankey (version 0.0.9).

### Sample Preparation and MALDI-MS Imaging

Wheat grain at DAP 4 and DAP 8 were harvested. The grains were then embedded with 2% CMC in the embedding box and frozen in a dry ice-ethanol bath. After the embedding medium has fully solidified and turned white, the box can be taken out and stored at - 80 °C. Before the section, samples were placed in a - 20 °C refrigerator to equilibrate for 1 h and fixed in three drops of distilled water for cutting. The seeds were sectioned at 20 μm thickness using a Leica CM1950 cryostat (Leica Microsystems GmbH, Germany) at - 20 °C. Subsequently, the seed sections were placed in groups on electrically conductive slides coated with indium tin oxide (ITO), and dried in a vacuum desiccator for 30 min. Desiccated sections mounted on ITO glass slides were sprayed using an HTX TM sprayer (Bruker Daltonics, Germany) with 15 mg/mL DHB, dissolved in 90%: 10% acetonitrile: water. The sprayer temperature was set to 70 °C, with a flow rate of 0.1 mL/min, and a pressure of 6 psi. 28 passes of the matrix were applied to slides with 5 s of drying time between each pass.

MALDI timsTOF MS imaging experiments were performed on a prototype Bruker timsTOF flex MS system (Bruker Daltonics, Bremen, Germany) equipped with a 10 kHz smartbeam 3D laser. Laser power was set to 80% and then fixed throughout the whole experiment. The mass spectra were acquired in positive mode, and the data were obtained over a mass range from m/z (mass-to-charge ratio) 50-1300 Da. Metabolites in the samples were identified by comparing accurate m/z values, retention times (RT), and fragmentation patterns. The imaging spatial resolution was set to 50 μm, and each spectrum consisted of 400 laser shots. MALDI-MS were normalized with the Root Mean Square, and the signal intensity in each image was shown as the normalized intensity. MS/MS fragmentations performed on the timsTOF flex MS system in the MS/MS mode were used for further detailed structural confirmation of the identified metabolites.

### Pseudotime analysis

To explore the molecular mechanism of cell differentiation and cell fate determination, we performed pseudo-time analysis using Monocle (v.2.20.0) (Qiu et al., 2017). Firstly, marker genes were identified using the differentialGeneTest function, employing the fullModelFormulaStr and reducedModelFormulaStr parameters to define differential expression while concurrently controlling for batch effects. Secondly, we set up the parameter ‘‘max_components” as two and method as “DDR Tree” to reduce the dimensionality of the data into two components. In the lower dimensional space, the “orderCells” function was performed to order the cells in pseudo-time according to the transcriptome correlation. Further, we performed the “plot_cell_trajectory” function to visualize the cell trajectory. To designate the priori “start point” in the trajectory, the “orderCells” function was performed again with appointing the “root_state”. The branch-dependent genes were analyzed by “BEAM” function. Then, we performed “plot_genes_branched_heatmap” to demonstrate the bifurcation of gene expression along two branches.

### In situ hybridization

In situ hybridization experiments and the detection of hybridized signals were carried out as described by Meng et al (Meng et al., 2017). Wheat grain was fixed in 10% formaldehyde overnight at 4°C. Then specimens were embedded in paraplast (Sigma, USA) and sectioned at 8.0 μm. Antisense and sense RNA probes were synthesized using a digoxigenin RNA labeling kit (Sigma-Aldrich, USA). Antisense RNA probe for *TaJEKYLL* genes was amplified by PCR using the upper primer 5′-ATGGCGGCTCGCACTG - 3′ and lower primer 5′ - GAATTGTAATACGACTCACTATAGGGTGGAGCTGCATCCTCTTCA - 3′. Sense RNA probe was amplified by PCR using the upper primer 5′-GAATTGTAATACGACTCACTATAGGGATGGCGGCTCGCACTG - 3′ and lower primer 5′ - TGGAGCTGCATCCTCTTCA - 3′.

### Subcellular localization

For subcellular localization, we transferred a 35S::TaMADS58-A1-GFP construct into the pMDC83 vector and then introduced it into into the protoplasts of wheat leaf.

To observe the localization of TaMADS58-A1, the coding sequences of *OsH2B* and *TaMADS58-A1* were amplified and ligated into the pCsVMV-mCherry-N-1300 and pCAMBIA1301-GFP vectors to generate CsVMV::H2B:mCherry and 35S::TaMADS58-A1:GFP, respectively, following the previously described protocol for N. benthamiana leaves (Liu et al., 2020). Similarly, colocalization of TaMADS58-A1 with the H2B:mCherry was observed according to a previous report (Zhang et al., 2025a). The fluorescence was visualized with a confocal scanning microscope (Nikon A1 HD25) 48-72 h after infiltration.

### Plasmid construction and wheat transformation

For wheat transformation, *TaMADS58-A1* and *TaJEKYLL-A1* full length cDNA was inserted into the Gateway-compatible OE vector PC186 (pUbi:: GWOE::Nos). The 2096-2200-bp region (counted from the start codon) of *TaJEKYLL-A1*, which was identified to be specific in wheat genome and conserved in *TaJEKYLLs*, was cloned into the amiR vector PC186. The PCR primers were listed in Table S10. The CRISPR-*TaJEKYLLs* and *TaMADS58-A1-OE* vectors were introduced into hexaploid wheat (cv. Fielder) via Agrobacterium-mediated transformation as described by Ishida et al48. PCR, herbicide (glufosinate) spraying and a QuickStix Kit for bar were used to verify the positive transgenic plants.

### RNA extraction, RT-qPCR analysis, and RNA-Seq

Total RNAs were extracted from grain at 4 DAP dand 8 DAP of Fielder, *TaMADS58-A1-OE*-1, *TaMADS58-A1-OE*-2, *TaMADS58-A1-OE*-3, *TaJEKYLLs*-KO-1, *TaJEKYLLs*-KO-2 and *TaJEKYLLs*-KO-3 using the FastPure Plant Total RNA Isolation Kit (AC0305, SPARKeasy) according to the manufacturer’s protocol. First-strand cDNAs were generated using a reverse transcription kit (AG0305-B, SPARKeasy). Subsequent RT-qPCR assays were performed using the SYBR Green PCR Master Mix (Q121-02/03, Vazyme Biotech). with each RT-qPCR assay being replicated at least three biological replicates and three technical replicates. Wheat *Actin* gene (*TaActin*, *TraesCS5A02G124300*) was used as an internal reference. Relevant primer sequences are given in Supplemental Data Set S11.

### Bulk RNA-seq analysis

RNA-seq libraries’ construction and sequencing platform were the same as previous description (Zhao *et al*., 2023), by Annoroad Gene Technology. To quantify the number of paired reads mapped to each gene, featureCounts (version 2.0.1) with the parameter “-p” was utilized (Ramírez et al., 2014). The resulting counts files were then used for differential expression analysis using DESeq2 (version 1.26.0) (Love et al., 2014), with a threshold of “|Log2 Fold Change| ≥ 0.75 and FDR ≤ 0.05” to identify differentially expressed genes. The raw counts were normalized to TPM (Transcripts Per Kilobase Million) for gene expression quantification.

### Gene co-expression network analysis

hdWGCNA was applied to high-dimensional single-cell RNA-seq data using the R package hdWGCNA (version 0.4.06) (Morabito *et al*., 2023). Genes expressed in ≥5% of cells were selected via the SetupForWGCNA function. The k-nearest neighbors (KNN) algorithm aggregated similar cells to generate a meta-cell gene expression matrix. Parameter scanning and soft threshold determination were performed using the TestSoftPowers and ConstructNetwork functions, respectively. Module eigengenes (MEs) summarized co-expression module profiles, and gene connectivity (kME) was quantified using the ModuleConnectivity function. Genes with high kME values were identified as hub genes.

### Microscopy and cell length measurement

The developing grains were collected from *TaJEKYLLs*-KO, *TaMADS58-A1-OE*, *tajekyllaa*, *tamasd58aa* and WT plants, grains were prepared: soaked in a 75% optimal cutting temperature compound (OCT, Sakura Finetek Europe B.V.) solution under vacuum for 5 minutes, then embedded in OCT and stored at -80 ℃. Once equilibrated to -20 ℃, samples were sliced into 10 μm thick sections. The sections were imaged under a SteREO DiscoveryV20 microscope (Carl Zeiss). To measure the cell length, approximately 100 cells were analysed in each of at least six different grain sections using the ImageJ software.

### Dual-Luciferase assay

The reporter plasmids were generated by inserting promoter sequences into the cloning site upstream of the LUC CDS in the vector pGreen0080. Mutations of the promoter sequence were created using PCR amplification and validated by sequencing. The primers used are listed in Supplemental Data Set S11. The effector plasmids were prepared by inserting the CDS into the pUC18 vector containing the 35S promoter, total protein was extracted and analyzed on a luminometer (Promega 20/20). A commercial LUC analysis kit was used according to the manufacturer’s protocols (Promega). Three biological replicates were performed for each experiment.

### Electrophoretic Mobility Shift Assay

Recombinant protein purification and EMSA The full-length CDSs of *TaMADS58-A1* and *TaMADS29-A1* were inserted into the cloning sites of the pmal-c2xMBP expression vector. Recombinant proteins were produced in Escherichia coli DE3 (BL21) cells. Protein production was induced for 16 h by the addition of isopropyl b-d-1-thiogalactopyranoside (IPTG) at a final concentration of 0.625 mM when OD600 of the cells reached 0.6. The cells were then sonicated and the supernatants containing the fusion proteins were purified with MBP beads (Beyotime). Oligonucleotide probes were synthesized and labeled with biotin at the 5-end. EMSA were performed according to the LightShift Chemiluminescent EMSA Kit (Thermo). All oligonucleotides and primers are listed in Supplemental Data Set S11.

### Yeast one-hybrid assay

Yeast one-hybrid assays were performed according to the Yeastmaker Yeast Transformation System 2 User Manual (Clontech 630439). Briefly, the bait sequences were synthesized, cloned into pAbAi vector, and integrated into the genome of the Y1HGold yeast strain. The prey sequences were amplified, cloned into the pGADT7 vector, and fused with the bait yeast strain. The primers used are listed in Supplemental Data Set S11. To determine protein and DNA interaction, the bait yeast strains were selected on medium lacking uracil with different concentrations of the Aureobasidin A (AbA) antibiotic.

### Haplotype analysis

Natural variations identified from population transcriptomic data (Zhao *et al*., 2024) were employed to assess allelic and expression variations of *TaMADS58-B1* and *TaJEKYLL-A1.* Additionally, re-sequencing data of the Watkins collection (Cheng *et al*., 2024) were analyzed to examine natural variations within the coding regions, 2-kb promoter regions, and 2-kb downstream regions of these genes. Polymorphic sites meeting the following criteria were retained for subsequent analysis: missing rate < 0.5, minor allele frequency < 0.05, and heterozygosity < 0.5. Gene haplotypes were constructed using the geneHapR (version 1.2.4) package in R (Zhang et al., 2023), followed by statistical tests to examine phenotypic and expression differences among distinct haplotypes.

## Supporting information

Supplemental Data Set

## Ethics approval and consent to participate

Not applicable.

## Consent for publication

Not applicable.

## Availability of data and materials

The raw sequencing data of SnRNAseq and bulk RNA-seq are available at Genome Sequence Archive of the National Genomics Data Center, China National Center (PRJCA038633). The spatial transcriptomics during seed development was download from NCBI (https://www.ncbi.nlm.nih.gov/) under the BioSample of SAMN41774528. RNA-seq and ATAC-seq during embryo development were download from the Genome Sequence Archive (GSA) under accession number PRJCA008382. RNA-seq and ATAC-seq generated during endosperm development were download from GSA under accession number PRJCA022666. All scripts used in this work are available at Github (https://github.com/ZhangZhaoheng24/snRNA_wheat_seed).

## Competing interests

The authors declare that they have no competing interests.

## Funding

This research is supported by the Natural Science Foundation of Shandong Province (SYS202206), National Natural Science Foundation of China (U24A20389), National Key Research and Development Program of China (2023YFF1000602), Key R&D Program of Shandong Province, China (2024CXPT072), the Key Science and Technology Project of Xinjiang Production and Construction Corps (2024AB007).

## Author contributions

Y.C. and J.X. conceived and supervised the study. Z.Z. performed the bioinformatics analyses. X.L., Q.Z. and F.Z. carried out the experimental work. X.-L.L. prepared the snRNA-seq libraries. Z.Z. and X.L. prepared the figures. Y.C., Z.Z. and X.L. wrote the manuscript, and X.L. and J.X. revised it. All authors discussed the results and approved the final manuscript.

## Acknowledgements

We thank Ms. Yanyan Liang of Peking University Institute of Advanced Agricultural Sciences for providing the wheat genetic transformation.

## Additional information

Fig. S1 Summary of the wheat grain snRNA-seq

A. Overview of the snRNA-seq and cluster annotation workflow using snRNA-seq libraries generated from DAP 4 and 8

B-D. SnRNA-seq quality metrics showing the distribution of UMI counts (A), gene counts (B) and cell counts (C) across snRNA-seq libraries.

E. Bar plot showing the cell count across cell cluster. The number of cell genes for each cell cluster is given on the top.

F-G. Violin plot showing gene count (E) and UMI counts (F) across cell cluster.

H Dotplot showing the expression pattern of the selected putative cell-type marker genes. The circle size indicates the relative percentage of cells expressing the marker genes, and the color represents the relative expression level.

I UMAP plots of marker genes revealing the identities of cell clusters (Embryo and Scutellum). Each dot represents a single cell, color-coded by expression level. Color scale represents normalized expression levels.

K UMAP plot showing the enrichment of cell cycle phase specific genes. Color scale represents enrichment levels.

J. Spatial visualization of the unbiased spot clustering for DAP 4 and DAP 8 sections. Merged bright field images and spatial clusters of the other three sections. The tissue/cell-type identity of each cluster was assigned based on the location of each cluster.

L. Heatmap showing Pearson correlation coefficient (PCC) across 11 cell-types.

Fig. S2 Integrative analysis of bulk RNA-seq, ATAC-seq, snRNA-seq, and spatial metabolomics during early wheat grain development

A. K-mean clustering of differentially expressed genes during grain development based on bulk RNA-seq (left). UMAP visualization showing the enrichment patterns of genes from each bulk RNA-seq cluster across the cell clusters in the snRNA-seq (right).

B. K-mean clustering of genes with differentially chromatin accessibility during grain development based on in bulk ATAC-seq (left). UMAP visualization showing the enrichment patterns of genes from each bulk ATAC-seq cluster across the cell clusters in the snRNA-seq (right).

C. Schematic diagram of the differentiation and segmentation of the spatial map of hormone metabolism

D. UMAP plot shown the expression enrichment levels of auxin related biosynthetic, metabolic and signaling genes. Color-coded based on the expression levels of the selected gene sets.

E. The MALDI-MS images of the 3-Indoleacrylic acid in Fielder grain at DAP 4 and DAP 8. The boxplot shown the quantified signal intensities across defined seed regions.

F. UMAP plot shown the expression enrichment levels of abscisic acid and brassinosteroids related biosynthetic, metabolic and signaling genes. Color-coded based on the expression levels of the selected gene sets.

G. UMAP plot show the expression levels of *TaGlu-B5* between DAP 4 and DAP 8.

H. UMAP plot show the expression levels of *TaBEIIa-A1* between DAP 4 and DAP 8.

Fig. S3 Developmental trajectories of wheat early endosperm and branch-specific gene

A. Developmental trajectory of seed cells split by cell types.

B. Scatter plot of grain size-related traits (GW, grain weight, GL, grain length, SDS, SDS sedimentation value) against the expression level values of TaMADS58-A1/B1/D1 in previously published population transcriptomes.

TaMADS58 regulates pericarp development while negatively modulating grain size and enhancing wheat end-use quality.

Fig. S4 TaMADS58 regulates the thickness of the pericarp of wheat grains and seed storage protein accumulation.

A. Expression pattern of TaMADS59s in grain at DAP 10 as detected by RNA in situ hybridization. P, pericarp, En, endosperm, Em, embryo, AL, aleurone layer. Scale bars, 200 μm.

B. Grain traits of the tamads58-b1. The grains exhibited significant differences in length and width. N = 10 grains, Bars = 1 cm.

C. Quantification of grain agronomic traits related traits between the WT plants and TaMADS58-A1-OE lines. Student’s t test was used to determine the difference significance between TaMADS58-A1-OE and WT. *, p ≤ 0.05, **, p ≤ 0.01, Data represent mean ± SD (n = 15 biological replicates).

D. The localization of TaMADS58-A1 was achieved by infecting N. benthamiana leaves with A. tumefaciens strains containing TaMADS58-A1. Fluorescence signals were detected at 48 hpi. Bar, 50 µm.

E. Sample similarity analysis between Fielder and TaMADS58-A1-OE lines.

F. GO enrichment for up- and down-regulated genes in TaMADS58-A1-OE line. False discovery rate (FDR) by Benjamini-Hochberg procedure.

G. Y1H assay of TaMADS58-A1 binds to TaLWM-GS-D1 and TaCDPK1-A1 promoter.

Fig. S5 Single-cell co-expression networks

A. UMAP plot showing the expression distribution of hub genes for each module across the 11 clusters.

B. UMAP heatmap showing the relative expression levels of SWEET, SUC, STP, NMG, AtUmamiT and AAAP family genes.

Fig. S6 TaJEKYLL mutations repressed the nutrient transportation and starch biosynthesis in early developing wheat grains.

A. Grain traits of the tajekyll-a1. The grains of different lines exhibited significant differences in length and width. N = 10 grains, Bars = 1 cm.

B. Transverse section of Developing grains revealed the nucellar projection degradation in the tajekyll-a1. Red boxes in the top panel were magnified and represented in the lower panel. Bars=200 μm.

C. Statistic comparison of nucellar projection width and area between KN9204 and tajekyll-a1. n = 3. Significance was determined using Student’s t-test, **P < 0.01.

D. Schematic of TaJEKYLL homeolog genomic structures showing the target sites and PAMs for single guide RNAs used for mutagenesis by CRISPR-Cas9 in the Fielder background. Sequences of Fielder and three recovered mutant alleles (designated KO1, KO2 and KO3) are shown (bottom).

E. Expression analysis of tajekyll-abD and its downstream target genes in WT and KO1 lines. TaACTIN was used as the internal control. Data are presented as means ± SD. P-values were determined by Student’s t-test. n = 3.

Fig. S7 Haplotype analysis of TaMADS58-B1 and TaJEKYLL-A1 in Watkins population.

A. Schematic showing the polymorphism for each haplotype of TaMADS58-B1. The coordinate is related to the transcription start site (TSS).

B. The SNPs and haplotypes of TaMADS58-B1.

C. The percentage of accessions carrying different haplotypes of TaMADS58-B1 in each ecological zone worldwide. The size of pie charts in the geographical map shows the number of accessions, with percentages of the three alleles in different colors.

D. Schematic showing the polymorphism for each haplotype of TaJEKYLL-A1. The coordinate is related to the transcription start site (TSS).

E. The SNPs and haplotypes of TaJEKYLL-A1.

F. The percentage of accessions carrying different haplotypes of TaJEKYLL-A1 in each ecological zone worldwide. The size of pie charts in the geographical map shows the number of accessions, with percentages of the three alleles in different colors.

## Supplemental Data Set

Supplemental Data Set S1. List of marker genes for 11 cell clusters identified from snRNA-seq and spatial transcriptome.

Supplemental Data Set S2. List of cell cycle-related genes

Supplemental Data Set S3. The comparison of marker genes generated from wheat and maize endosperm.

Supplemental Data Set S4. Gene expression bias based on snRNA-seq and bulk RNA-seq in seed.

Supplemental Data Set S5. Examples of differenfially expressed genes between DAP 4 and DAP 8 within each cell type.

Supplemental Data Set S6. The sources of all hormone genes used in this study. Supplemental Data Set S7. The TF family of branch-specific expressed TF in differentiation trajectories.

Supplemental Data Set S8. The expression levels of Differentially expressed SSP and starch synthesis genes between Fielder and *TaMADS58-A1*-OE.

Supplemental Data Set S9. Examples of hWGCNA module specific-genes.36 Supplemental Data Set S10. The C/N transporter genes used in this study.

Supplemental Data Set S11. The primers and oligonucleotides used in this study.

## Notes

### Competing Interest Statement

The authors have declared no competing interest.

## References

Albacete, A.A., Martínez-Andújar, C., and Pérez-Alfocea, F. (2014). Hormonal and metabolic regulation of source-sink relations under salinity and drought: from plant survival to crop yield stability. Biotechnology advances 32:12–30. 10.1016/j.biotechadv.2013.10.005.

Andreatta, M., and Carmona, S.J. (2021). UCell: Robust and scalable single-cell gene signature scoring. Comput. Struct. Biotechnol. J. 19:3796–3798. 10.1016/j.csbj.2021.06.043.

Calderini, D.F., Castillo, F.M., Arenas, M.A., Molero, G., Reynolds, M.P., Craze, M., Bowden, S., Milner, M.J., Wallington, E.J., Dowle, A., et al. (2021). Overcoming the trade-off between grain weight and number in wheat by the ectopic expression of expansin in developing seeds leads to increased yield potential. New Phytol 230:629–640. 10.1111/nph.17048.

Chen, Y., Song, W., Xie, X., Wang, Z., Guan, P., Peng, H., Jiao, Y., Ni, Z., Sun, Q., and Guo, W. (2020). A Collinearity-Incorporating Homology Inference Strategy for Connecting Emerging Assemblies in the Triticeae Tribe as a Pilot Practice in the Plant Pangenomic Era. Mol Plant 13:1694–1708. 10.1016/j.molp.2020.09.019.

Cheng, S., Feng, C., Wingen, L.U., Cheng, H., Riche, A.B., Jiang, M., Leverington-Waite, M., Huang, Z., Collier, S., Orford, S., et al. (2024). Harnessing landrace diversity empowers wheat breeding. Nature 632:823–831. 10.1038/s41586-024-07682-9.

Domínguez, F., and Cejudo, F.J. (2014). Programmed cell death (PCD): an essential process of cereal seed development and germination. Front Plant Sci 5:366. 10.3389/fpls.2014.00366.

Domínguez, F., Moreno, J., and Cejudo, F.J. (2001). The nucellus degenerates by a process of programmed cell death during the early stages of wheat grain development. Planta 213:352–360. 10.1007/s004250000517.

Feldman, M., Levy, A.A., Fahima, T., and Korol, A. (2012). Genomic asymmetry in allopolyploid plants: wheat as a model. J Exp Bot 63:5045–5059. 10.1093/jxb/ers192.

Feng, S., Liu, Z., Chen, H., Li, N., Yu, T., Zhou, R., Nie, F., Guo, D., Ma, X., and Song, X. (2024). PHGD: An integrative and user-friendly database for plant hormone-related genes. Imeta 3:e164. 10.1002/imt2.164.

Fu, Y., Xiao, W., Tian, L., Guo, L., Ma, G., Ji, C., Huang, Y., Wang, H., Wu, X., Yang, T., et al. (2023). Spatial transcriptomics uncover sucrose post-phloem transport during maize kernel development. Nat Commun 14:7191. 10.1038/s41467-023-43006-7.

Gong, G., Jia, H., Tang, Y., Pei, H., Zhai, L., and Huang, J. (2024). Genetic analysis and QTL mapping for pericarp thickness in maize (Zea mays L.). BMC Plant Biol 24:338. 10.1186/s12870-024-05052-1.

Guo, X., Wang, Y., Zhao, C., Tan, C., Yan, W., Xiang, S., Zhang, D., Zhang, H., Zhang, M., Yang, L., et al. (2025). An Arabidopsis single-nucleus atlas decodes leaf senescence and nutrient allocation. Cell 188:2856–2871.e2816. 10.1016/j.cell.2025.03.024.

Hao, Y., Hao, S., Andersen-Nissen, E., Mauck, W.M., 3rd, Zheng, S., Butler, A., Lee, M.J., Wilk, A.J., Darby, C., Zager, M., et al. (2021). Integrated analysis of multimodal single-cell data. Cell 184:3573–3587.e3529. 10.1016/j.cell.2021.04.048.

Lee, T.A., Illouz-Eliaz, N., Nobori, T., Xu, J., Jow, B., Nery, J.R., and Ecker, J.R. (2025). A single-cell, spatial transcriptomic atlas of the Arabidopsis life cycle. Nat Plants 11:1960–1975. 10.1038/s41477-025-02072-z.

Li, X., Wan, Y., Wang, D., Li, X., Wu, J., Xiao, J., Chen, K., Han, X., and Chen, Y. (2025). Spatiotemporal transcriptomics reveals key gene regulation for grain yield and quality in wheat. Genome biology 26:93. 10.1186/s13059-025-03569-8.

Liang, H., Zhou, J., and Chen, C. (2025). The aleurone layer of cereal grains: Development, genetic regulation, and breeding applications. Plant communications 6:101283. 10.1016/j.xplc.2025.101283.

Liu, C., Wu, T., Fan, F., Liu, Y., Wu, L., Junkin, M., Wang, Z., Yu, Y., Wang, W., Wei, W., et al. (2019). A portable and cost-effective microfluidic system for massively parallel single-cell transcriptome profiling. bioRxiv:818450. 10.1101/818450.

Liu, G., Zhang, R., Li, S., Ullah, R., Yang, F., Wang, Z., Guo, W., You, M., Li, B., Xie, C., et al. (2023). TaMADS29 interacts with TaNF-YB1 to synergistically regulate early grain development in bread wheat. Science China. Life sciences 66:1647–1664. 10.1007/s11427-022-2286-0.

Liu, J., Zhu, Y., Xian, M., Shen, L., Li, Y., Song, J., and Dai, Z. (2022). Structural development of the nutrient transfer tissues in different waxy wheat grain. Cereal Research Communications 50:953–964. 10.1007/s42976-022-00249-2.

Liu, J., Zhu, Y., Liu, X., Song, J., Tang, L., Shen, L., and Dai, Z. (2025). Morphological development of the endosperm epidermal cells in waxy wheat cultivars. Protoplasma 262:957–977. 10.1007/s00709-025-02034-4.

Liu, Y., Hou, J., Wang, X., Li, T., Majeed, U., Hao, C., and Zhang, X. (2020). The NAC transcription factor NAC019-A1 is a negative regulator of starch synthesis in wheat developing endosperm. J Exp Bot 71:5794–5807. 10.1093/jxb/eraa333.

Love, M.I., Huber, W., and Anders, S. (2014). Moderated estimation of fold change and dispersion for RNA-seq data with DESeq2. Genome Biol 15:550. 10.1186/s13059-014-0550-8.

Meng, W.J., Cheng, Z.J., Sang, Y.L., Zhang, M.M., Rong, X.F., Wang, Z.W., Tang, Y.Y., and Zhang, X.S. (2017). Type-B ARABIDOPSIS RESPONSE REGULATORs Specify the Shoot Stem Cell Niche by Dual Regulation of WUSCHEL. Plant Cell 29:1357–1372. 10.1105/tpc.16.00640.

Millsteed, T., Kainer, D., Sullivan, R., Sun, X., Li, K.L., Mao, L., Macdonald, A., and Henry, R.J. (2025). Spatial Transcriptomics of Developing Wheat Seed Reveals Concentric Gene Expression Zones and Subgenome Biased Expression of Key Genes. Plant Biotechnol J 23:5934–5949. 10.1111/pbi.70351.

Morabito, S., Reese, F., Rahimzadeh, N., Miyoshi, E., and Swarup, V. (2023). hdWGCNA identifies co-expression networks in high-dimensional transcriptomics data. Cell Rep Methods 3:100498. 10.1016/j.crmeth.2023.100498.

Patrick, J.W., and Offler, C.E. (2001). Compartmentation of transport and transfer events in developing seeds. J Exp Bot 52:551–564.

Pei, H., Li, Y., Liu, Y., Liu, P., Zhang, J., Ren, X., and Lu, Z. (2023). Chromatin accessibility landscapes revealed the subgenome-divergent regulation networks during wheat grain development. aBIOTECH 4:8–19. 10.1007/s42994-023-00095-8.

Peukert, M., Thiel, J., Peshev, D., Weschke, W., Van den Ende, W., Mock, H.P., and Matros, A. (2014). Spatio-temporal dynamics of fructan metabolism in developing barley grains. Plant Cell 26:3728–3744. 10.1105/tpc.114.130211.

Picard, C.L., Povilus, R.A., Williams, B.P., and Gehring, M. (2021). Transcriptional and imprinting complexity in Arabidopsis seeds at single-nucleus resolution. Nat Plants 7:730–738. 10.1038/s41477-021-00922-0.

Qiu, X., Hill, A., Packer, J., Lin, D., Ma, Y.A., and Trapnell, C. (2017). Single-cell mRNA quantification and differential analysis with Census. Nat Methods 14:309–315. 10.1038/nmeth.4150.

Radchuk, V., Sharma, R., Potokina, E., Radchuk, R., Weier, D., Munz, E., Schreiber, M., Mascher, M., Stein, N., Wicker, T., et al. (2019). The highly divergent Jekyll genes, required for sexual reproduction, are lineage specific for the related grass tribes Triticeae and Bromeae. Plant J 98:961–974. 10.1111/tpj.14363.

Ramírez-González, R.H., Borrill, P., Lang, D., Harrington, S.A., Brinton, J., Venturini, L., Davey, M., Jacobs, J., van Ex, F., Pasha, A., et al. (2018). The transcriptional landscape of polyploid wheat. Science 361:eaar6089. doi:10.1126/science.aar6089.

Ramírez, F., Dündar, F., Diehl, S., Grüning, B.A., and Manke, T. (2014). deepTools: a flexible platform for exploring deep-sequencing data. Nucleic Acids Res 42:W187–191. 10.1093/nar/gku365.

Rasheed, A., Hao, Y., Xia, X., Khan, A., Xu, Y., Varshney, R.K., and He, Z. (2017). Crop Breeding Chips and Genotyping Platforms: Progress, Challenges, and Perspectives. Mol Plant 10:1047–1064. 10.1016/j.molp.2017.06.008.

Saada, S., Solomon, C.U., and Drea, S. (2021). Programmed Cell Death in Developing Brachypodium distachyon Grain. International journal of molecular sciences 2210.3390/ijms22169086.

Sajjad, M., Ma, X., Habibullah Khan, S., Shoaib, M., Song, Y., Yang, W., Zhang, A., and Liu, D. (2017). TaFlo2-A1, an ortholog of rice Flo2, is associated with thousand grain weight in bread wheat (Triticum aestivum L.). BMC Plant Biol 17:164. 10.1186/s12870-017-1114-3.

Sanchez-Bragado, R., Vicente, R., Molero, G., Serret, M.D., Maydup, M.L., and Araus, J.L. (2020). New avenues for increasing yield and stability in C3 cereals: exploring ear photosynthesis. Curr Opin Plant Biol 56:223–234. 10.1016/j.pbi.2020.01.001.

Sheraz, S., Wan, Y., Venter, E., Verma, S.K., Xiong, Q., Waites, J., Connorton, J.M., Shewry, P.R., Moore, K.L., and Balk, J. (2021). Subcellular dynamics studies of iron reveal how tissue-specific distribution patterns are established in developing wheat grains. New Phytol 231:1644–1657. 10.1111/nph.17440.

Wamalwa, M., Tadesse, Z., Muthui, L., Yao, N., Zegeye, H., Randhawa, M., Wanyera, R., Uauy, C., and Shorinola, O. (2020). Allelic diversity study of functional genes in East Africa bread wheat highlights opportunities for genetic improvement. Mol. Breed. 40:104. 10.1007/s11032-020-01185-x.

Wang, H.L., Offler, C.E., and Patrick, J.W. (1994). Nucellar projection transfer cells in the developing wheat grain. Protoplasma 182:39–52. 10.1007/BF01403687.

Wang, J., Ye, F., Chai, H., Jiang, Y., Wang, T., Ran, X., Xia, Q., Xu, Z., Fu, Y., Zhang, G., et al. (2025). Advances and applications in single-cell and spatial genomics. Science China. Life sciences 68:1226–1282. 10.1007/s11427-024-2770-x.

Wang, N., and Fisher, D.B. (1994). The Use of Fluorescent Tracers to Characterize the Post-Phloem Transport Pathway in Maternal Tissues of Developing Wheat Grains. Plant physiology 104:17–27. 10.1104/pp.104.1.17.

Wang, Y., Luo, Y., Guo, X., Li, Y., Yan, J., Shao, W., Wei, W., Wei, X., Yang, T., Chen, J., et al. (2024). A spatial transcriptome map of the developing maize ear. Nat Plants 10:815–827. 10.1038/s41477-024-01683-2.

Xiong, F., Yu, X.R., Zhou, L., Wang, F., and Xiong, A.S. (2013). Structural and physiological characterization during wheat pericarp development. Plant cell reports 32:1309–1320. 10.1007/s00299-013-1445-y.

Xurun, Y., Xinyu, C., Liang, Z., Jing, Z., Heng, Y., Shanshan, S., Fei, X., and Zhong, W. (2015). Structural development of wheat nutrient transfer tissues and their relationships with filial tissues development. Protoplasma 252:605–617. 10.1007/s00709-014-0706-0.

Yao, J., Marand, A.P., Bai, Y., Schmitz, R.J., and Fan, L. (2025). Advances in plant spatial multi-omics data analysis. Trends in plant science 10.1016/j.tplants.2025.10.005.

Yao, J., Chu, Q., Guo, X., Shao, W., Shang, N., Luo, K., Li, X., Chen, H., Cheng, Q., Mo, F., et al. (2024). Spatiotemporal transcriptomic landscape of rice embryonic cells during seed germination. Developmental cell 59:2320–2332.e2325. 10.1016/j.devcel.2024.05.016.

Yin, L.L., and Xue, H.W. (2012). The MADS29 transcription factor regulates the degradation of the nucellus and the nucellar projection during rice seed development. Plant Cell 24:1049–1065. 10.1105/tpc.111.094854.

Young, T.E., and Gallie, D.R. (2000). Programmed cell death during endosperm development. Plant molecular biology 44:283–301. 10.1023/a:1026588408152.

Yuan, Y., Huo, Q., Zhang, Z., Wang, Q., Wang, J., Chang, S., Cai, P., Song, K.M., Galbraith, D.W., Zhang, W., et al. (2024). Decoding the gene regulatory network of endosperm differentiation in maize. Nat Commun 15:34. 10.1038/s41467-023-44369-7.

Zhang, P., Gao, S., Jin, J., Li, H., Zhang, Y., Nie, X., Ma, T., Wang, H., Nadeem, B., Shi, P., et al. (2025a). Suppression of the Transcription Factor TaDOF7.6 Enhances Photosynthesis and Energy Metabolism to Boost Wheat Yield. Journal of agricultural and food chemistry 73:16660–16671. 10.1021/acs.jafc.5c03339.

Zhang, R., Jia, G., and Diao, X. (2023). geneHapR: an R package for gene haplotypic statistics and visualization. BMC Bioinf. 24:199. 10.1186/s12859-023-05318-9.

Zhang, W.H., Zhou, Y., Dibley, K.E., Tyerman, S.D., Furbank, R.T., and Patrick, J. W. (2007). Review: Nutrient loading of developing seeds. Functional plant biology : FPB 34:314–331. 10.1071/fp06271.

Zhang, X., Luo, Z., Marand, A.P., Yan, H., Jang, H., Bang, S., Mendieta, J.P., Minow, M.A.A., and Schmitz, R.J. (2025b). A spatially resolved multi-omic single-cell atlas of soybean development. Cell 188:550–567.e519. 10.1016/j.cell.2024.10.050.

Zhang, X., Wang, Y.P., Song, X., Zhou, L.Z., Yu, H., Yang, L., Wang, Y.K., Wang, X.Y., Wan, X.Y., Liu, Y., et al. (2025c). A Single-Cell-Resolution Spatial Transcriptomic Atlas Decodes Wheat Spike Development and Yield Potential. Mol Plant 10.1016/j.molp.2025.12.020.

Zhao, L., Yang, Y., Chen, J., Lin, X., Zhang, H., Wang, H., Wang, H., Bie, X., Jiang, J., Feng, X., et al. (2023). Dynamic chromatin regulatory programs during embryogenesis of hexaploid wheat. Genome biology 24:7. 10.1186/s13059-022-02844-2.

Zhao, L., Chen, J., Zhang, Z., Wu, W., Lin, X., Gao, M., Yang, Y., Zhao, P., Xu, S., Yang, C., et al. (2024). Deciphering the Transcriptional Regulatory Network Governing Starch and Storage Protein Biosynthesis in Wheat for Breeding Improvement. Advanced science (Weinheim, Baden-Wurttemberg, Germany) 11:e2401383. 10.1002/advs.202401383.

Zheng, Y., Fei, X., and Yu, X. (2017). Observation and Investigation of Starch Granules Within Wheat Pericarp and Endosperm. Agricultural Research 6:320–325. 10.1007/s40003-017-0270-x.

Zhu, M., Hsu, C.W., Peralta Ogorek, L.L., Taylor, I.W., La Cavera, S., Oliveira, D.M., Verma, L., Mehra, P., Mijar, M., Sadanandom, A., et al. (2025). Single-cell transcriptomics reveal how root tissues adapt to soil stress. Nature 642:721–729. 10.1038/s41586-025-08941-z.

Zhuo, J., Wang, K., Wang, N., Xing, C., Peng, D., Wang, X., Qu, G., Kang, C., Ye, X., Li, Y., et al. (2023). Pericarp starch metabolism is associated with caryopsis development and endosperm starch accumulation in common wheat. Plant science : an international journal of experimental plant biology 330:111622. 10.1016/j.plantsci.2023.111622.

