## Supplementary figures and images for "Single-cell transcriptomic landscapes reveal cell-type-specific regulatory mechanisms of nutrient accumulation and transport in wheat grain"

### Supplemental Figures

#### Slide 1
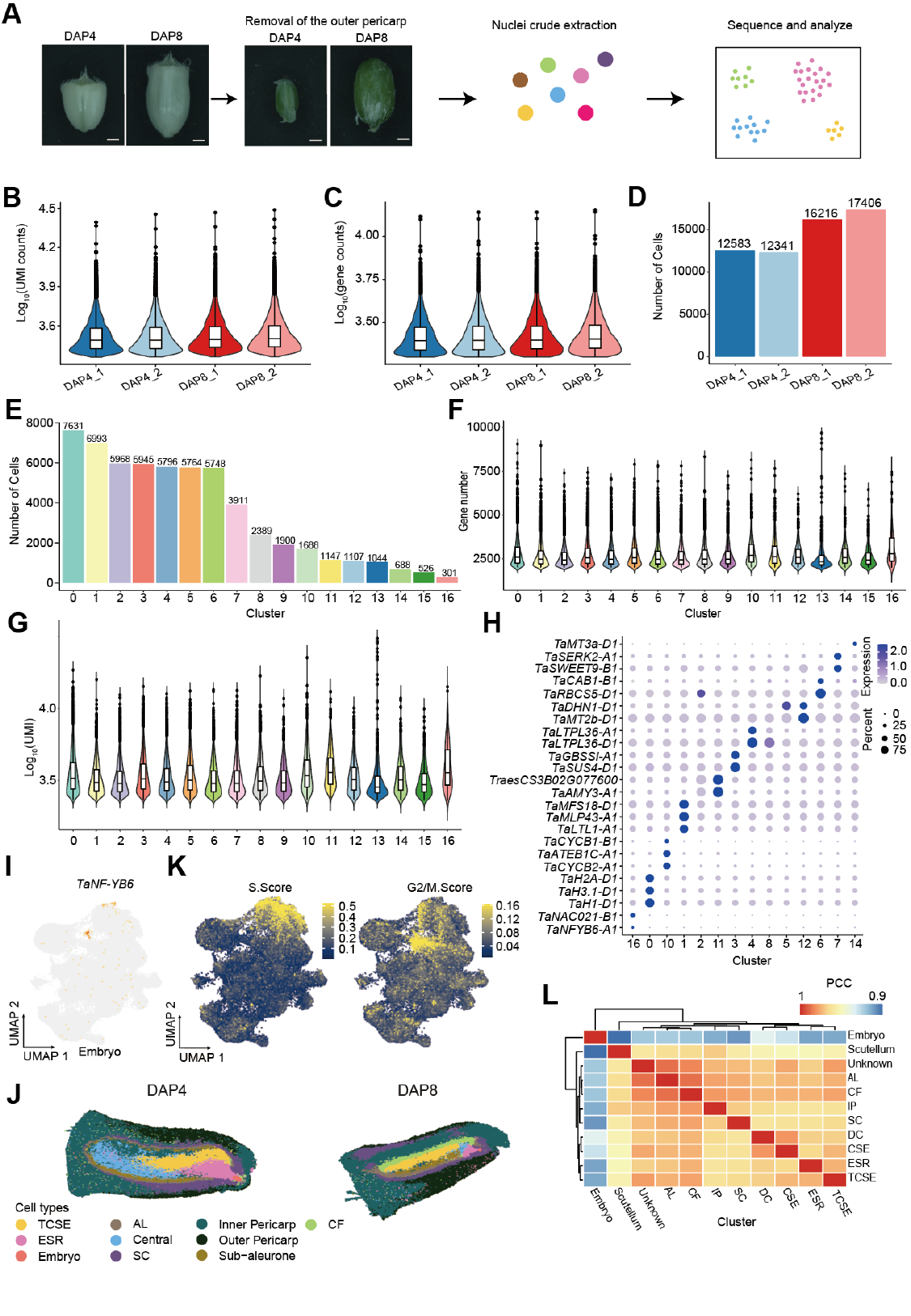

##### FigS1

#### Slide 2
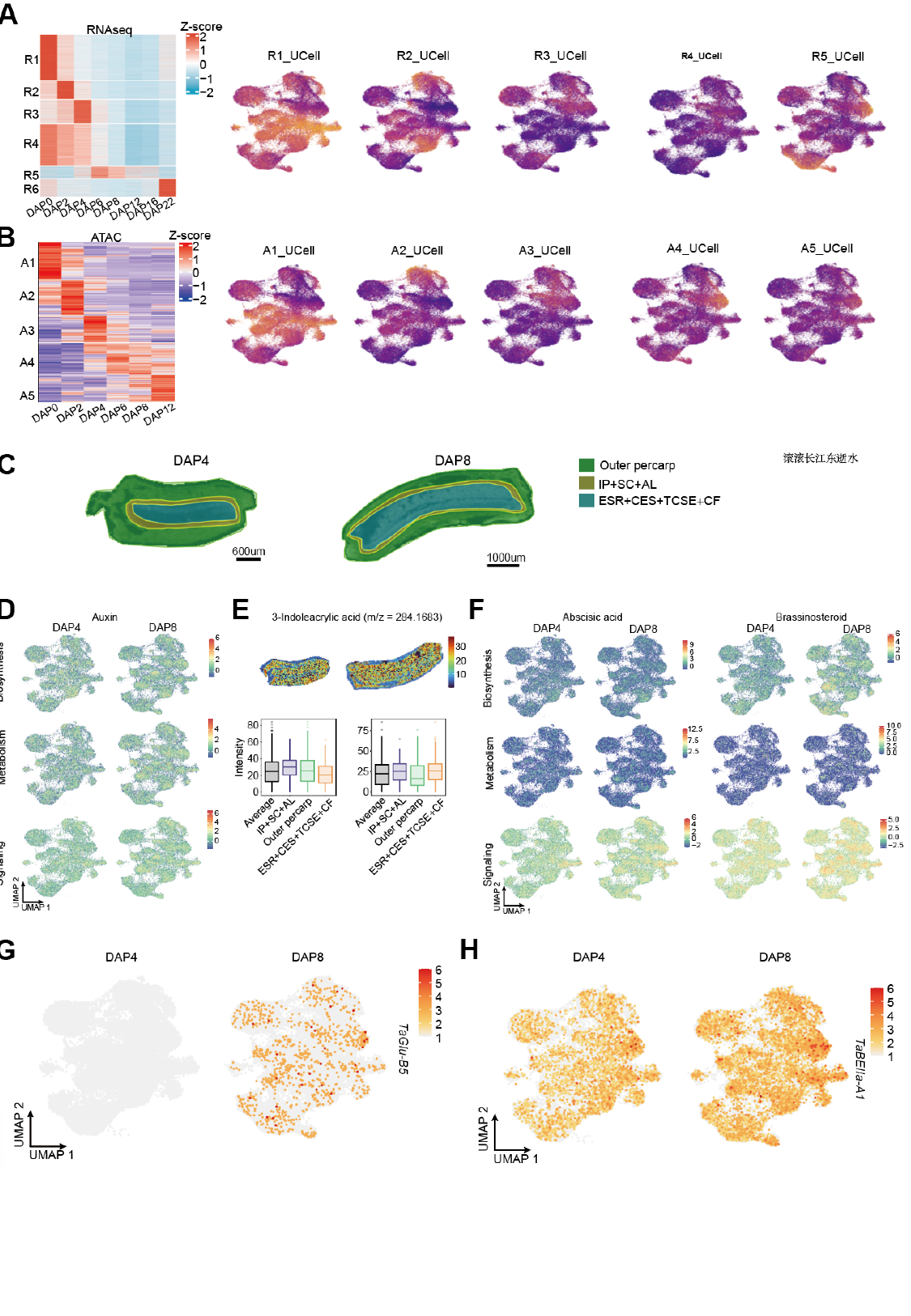

##### FigS2

#### Slide 3
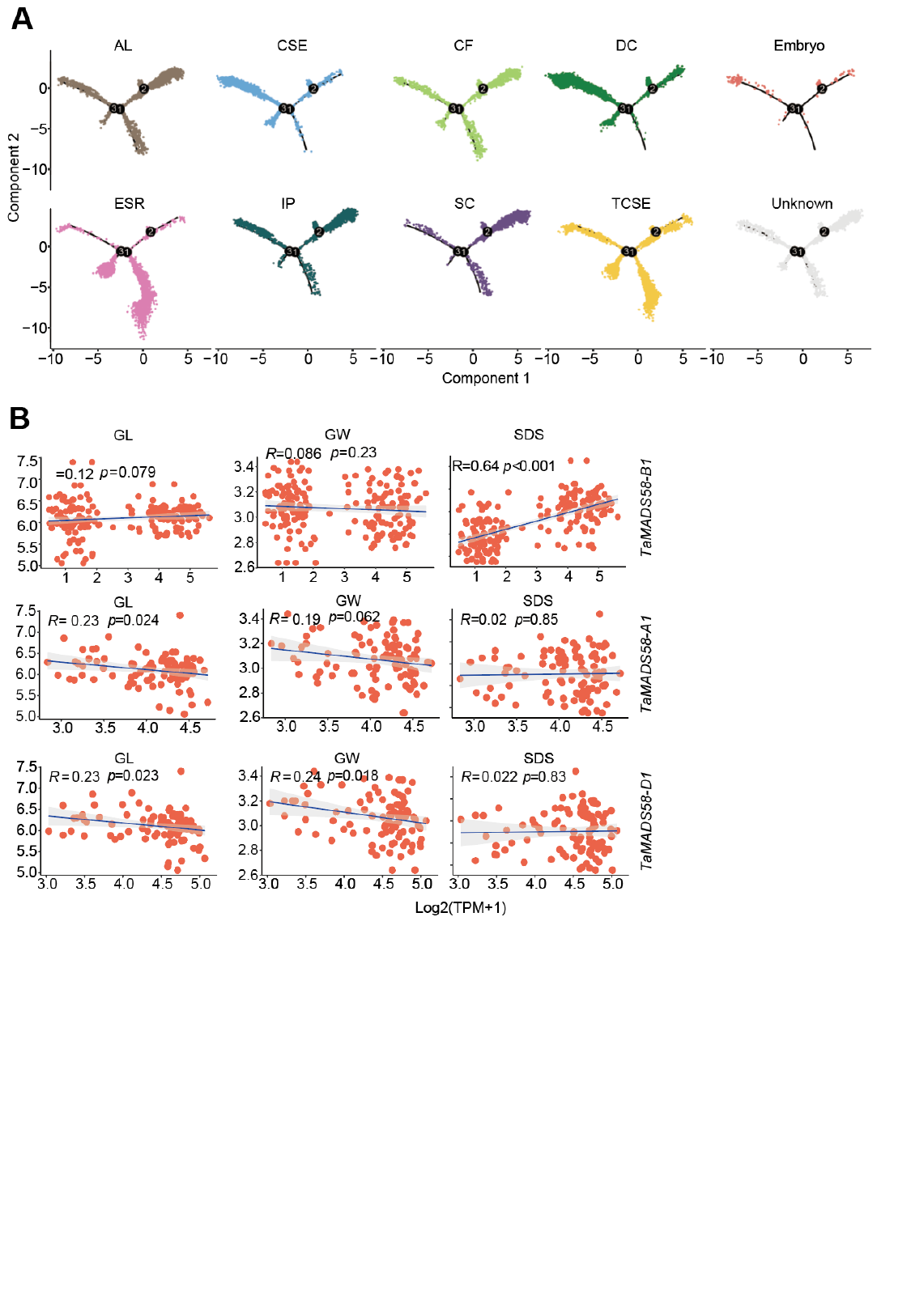

##### FigS3

#### Slide 4
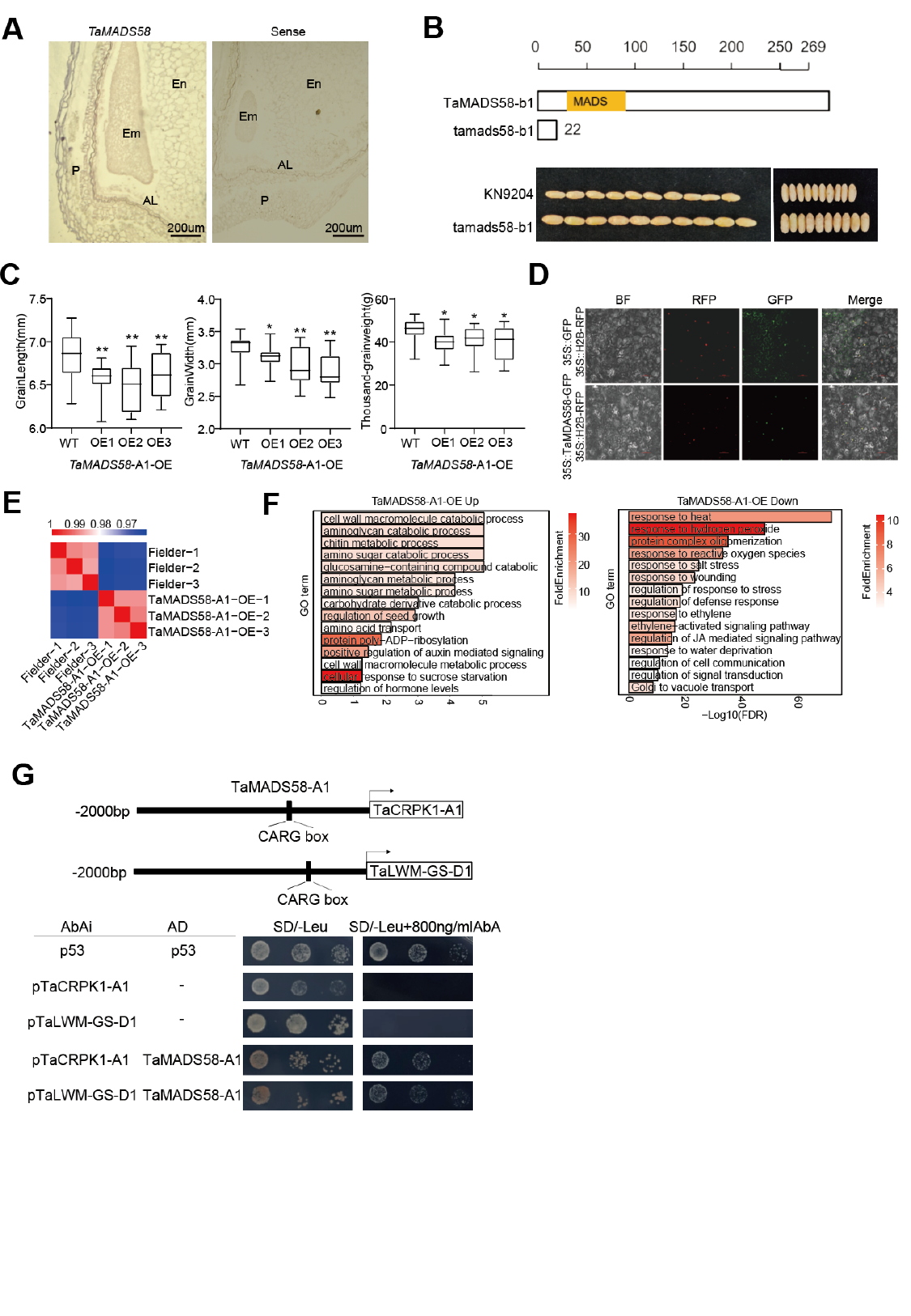

##### FigS4

#### Slide 5
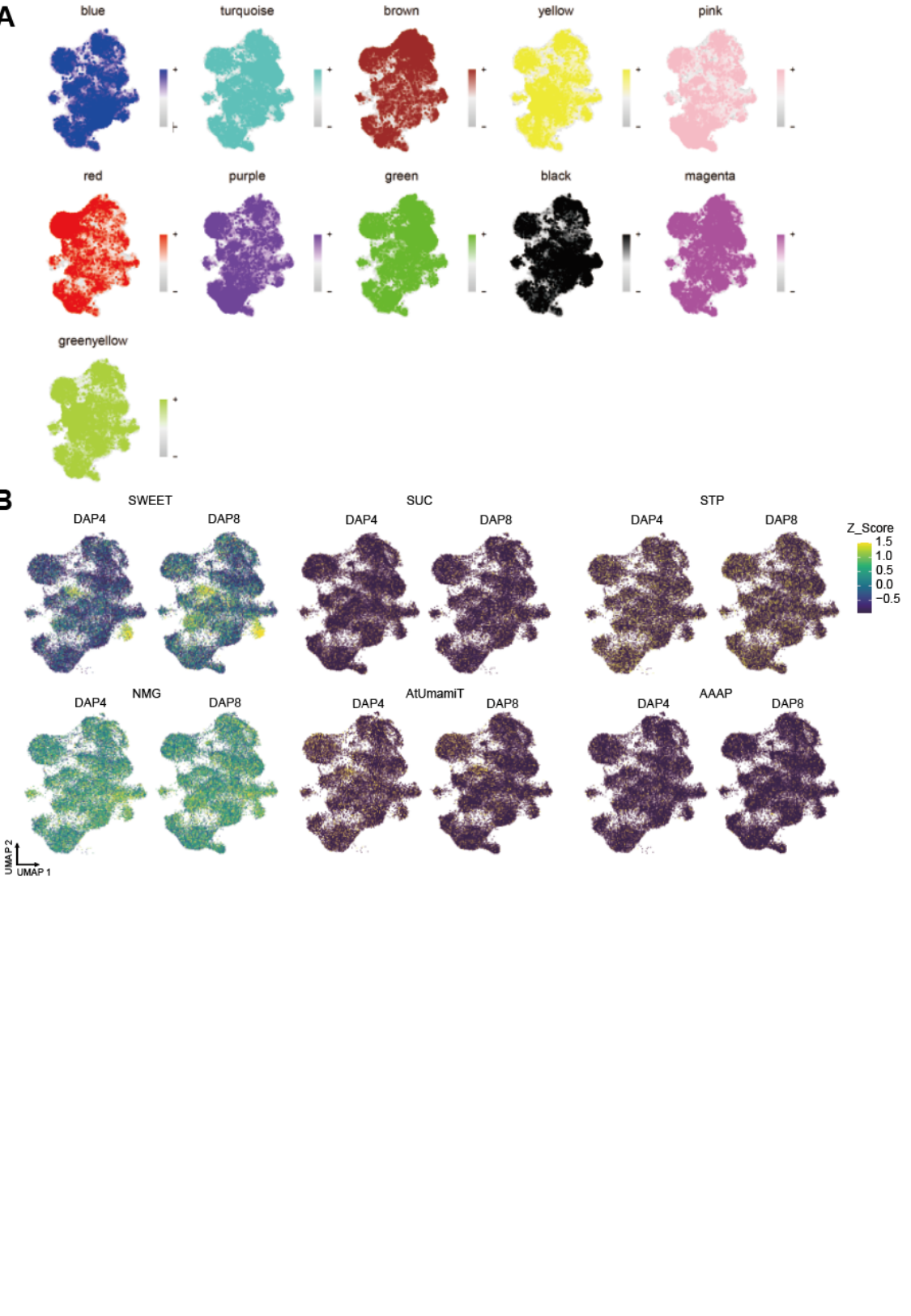

##### FigS5

#### Slide 6
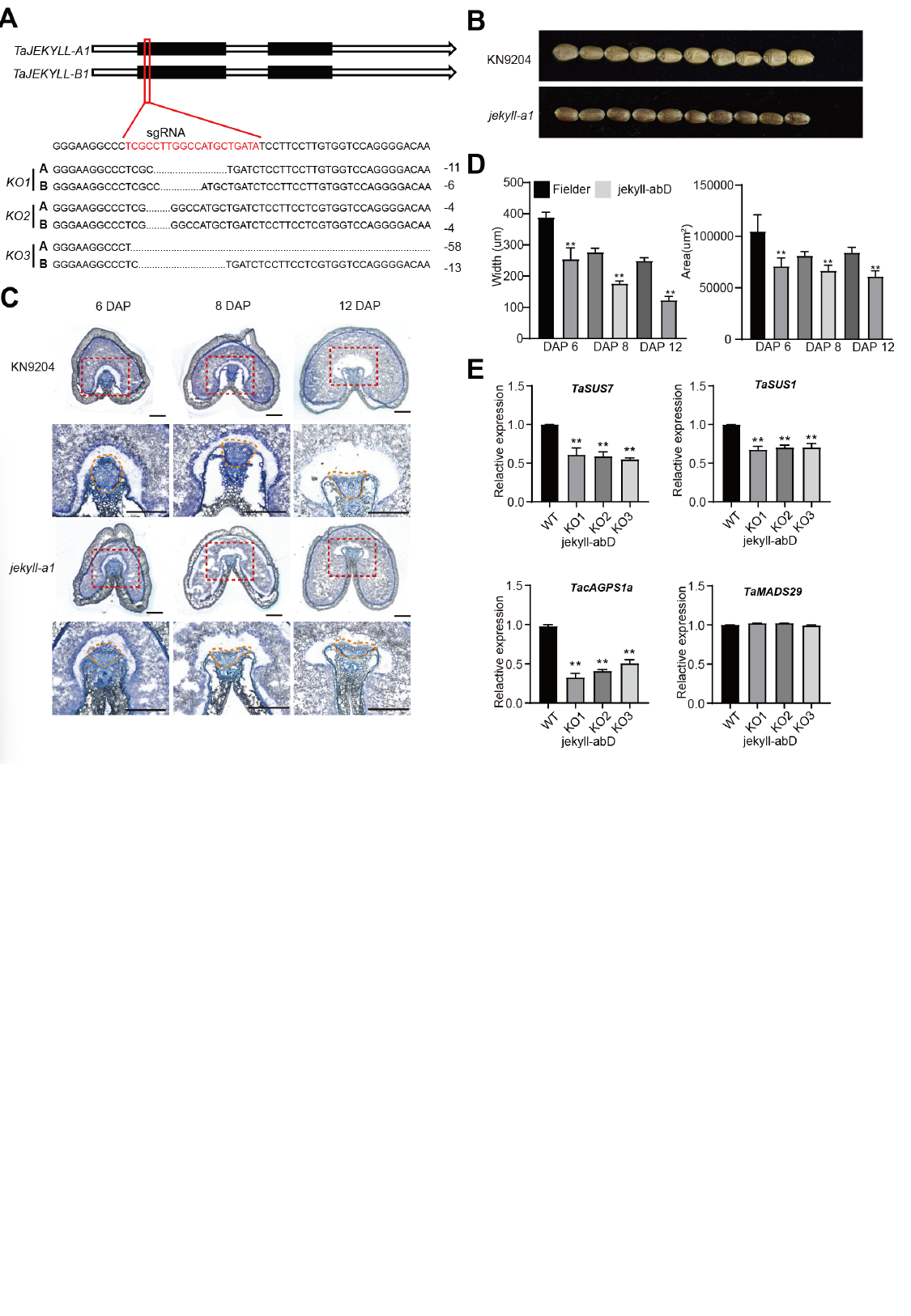

##### FigS6

#### Slide 7
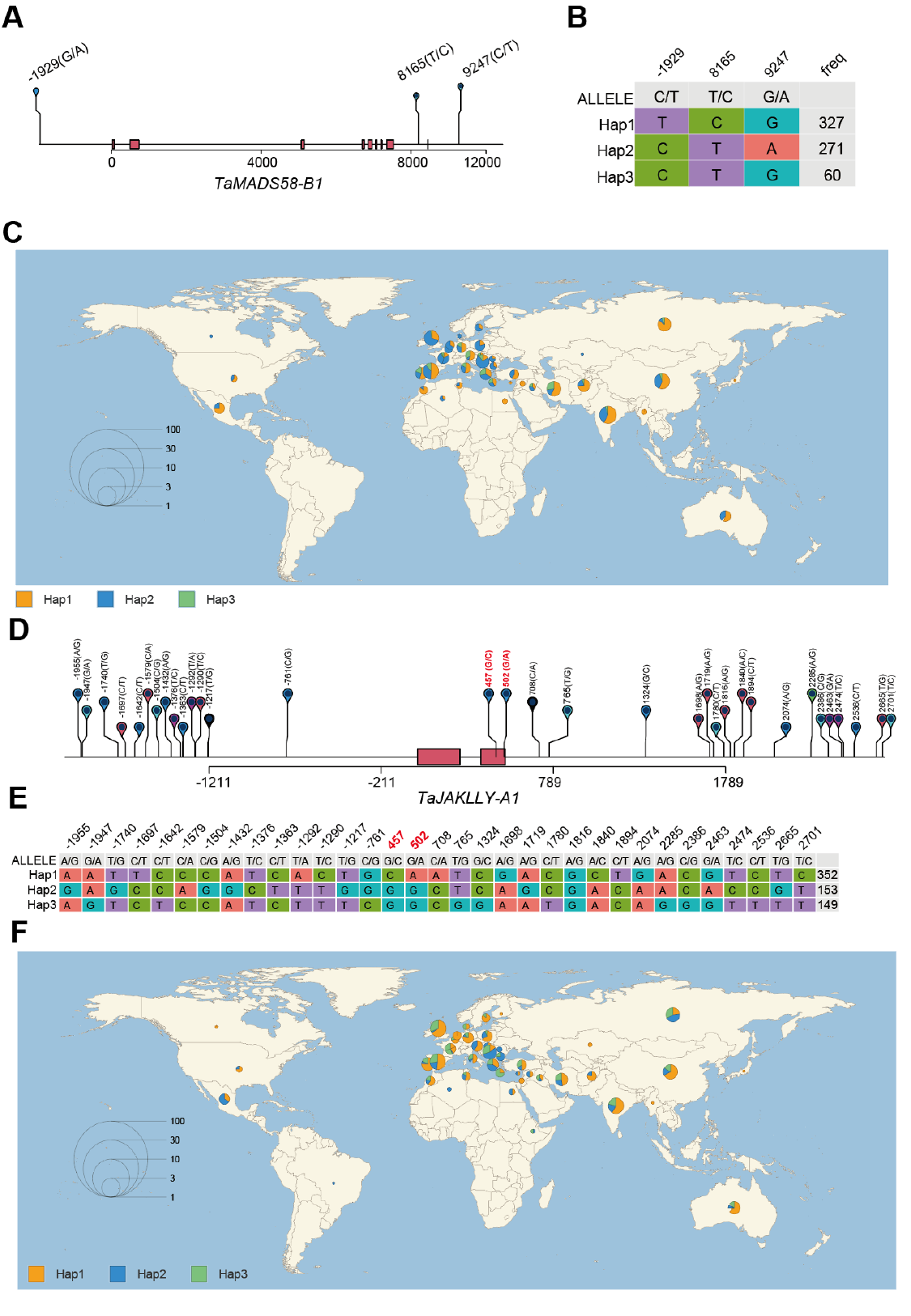

##### FigS7
